# High-resolution mapping and epistatic QTL of tomato fruit metabolism

**DOI:** 10.64898/2026.05.07.723420

**Authors:** Esra Karakas, Micha Wijesingha Ahchige, Dan Qian, Shai Torgeman, Björn Usadel, Dani Zamir, Alisdair R. Fernie, Saleh Alseekh

## Abstract

Tomato wild relatives are valuable genetic resources for trait discovery and understanding the genetic basis of fruit metabolism and quality. Yet, only a fraction of naturally occurring variation has been exploited. Here, we performed metabolite profiling of two large Backcross Inbred Line populations derived from crosses between the wild species *S. pennellii* accession LA5240 (Lost) and cultivated genotypes LEA (determinate) and TOP (indeterminate), including ∼1400 and ∼500 lines, respectively. High-resolution mapping identified enormous metabolic quantitative trait loci (mQTL), including a new locus on chromosome 12 associated with fruit sucrose accumulation that harbours *INVERTASE INHIBITOR* 3 (*SlINVINH3*) protein. Comparative analysis indicated that *SlINVINH3 is* highly expressed in wild *S. pennellii* 0716 fruit, whereas a six-amino acid deletion is present in its coding sequence compared with *S.pennellii* LA5240 and *S. lycopersicum*. We further demonstrated that in *SlINVINH3*-overexpressing tomato plants, only the *S. pennellii* LA5240 allele led to increased sucrose, accompanied by reduced fructose and glucose levels. Furthermore, the large population size enabled us to assess the epistatic interactions, with approximately 40% of interactions being more-than-additive and 60% less-than-additive. Our results demonstrate the power of permanent exotic populations to reveal hidden metabolic diversity and provide an approach for improving fruit quality through targeted breeding and metabolic engineering.

## Introduction

Exotic germplasm represents a rich source of genetic diversity that can be leveraged to improve crop traits (Zamir, 2001). Despite considerable successes in crop improvement, only a small fraction of naturally occurring variation has been exploited in breeding programs (Wang et al., 2017). The development of exotic library collections of marker-defined genomic segments introgressed from wild species into elite cultivars has enabled the discovery of numerous quantitative trait loci (QTL) and the identification of genes underlying agriculturally important traits, including stress adaptation and fruit quality (Collins et al., 2008).

A central yet often underappreciated feature of complex trait genetics is epistasis, the non-additive interaction between loci whereby phenotypic outcomes cannot be predicted from individual allelic effects alone. Epistatic interactions have been reported across diverse plant species and traits, from flowering time, yield and metabolic composition (Aguirre et al., 2023; Mackay, 2014; Soyk et al., 2017). By revealing masked genetic variation that remains hidden in single-locus analyses, epistasis reshapes our understanding of trait architecture and has important implications for breeding strategies and evolutionary adaptation (Napier et al., 2023).

Tomato (*Solanum lycopersicum*), has emerged as a powerful model for dissecting the genetic basis of quantitative traits, owing to its rich natural diversity and the availability of diverse mapping populations derived from wild relatives. These include introgression lines (ILs) (Eshed & Zamir, 1995), backcross inbred lines (BILs) (Ofner et al., 2016; Tanksley et al., 1996), recombinant inbred lines (RILs) (Paran et al., 1995), and MAGIC populations (Pascual et al., 2015). Among them, populations derived from *Solanum pennellii* have contributed substantially to our understanding of the genetic control of fruit morphology, yield and metabolism (Grandillo et al., 2007).

Primary metabolites such as sugars, organic acids and amino acids are key determinants of fruit quality and nutritional value. In tomato, sugar content strongly influences sweetness and consumer preference, making its genetic regulation a major breeding target (Klee & Tieman, 2018). Cultivated tomatoes predominantly accumulate the hexoses glucose and fructose, which correlate with soluble solid content and sweetness perception (Carrari & Fernie, 2006; Schauer et al., 2006). In contrast, wild species such as *S. pennellii* exhibit higher soluble solids and frequently accumulate sucrose, reflecting fundamental differences in carbohydrate metabolism and partitioning (Beckles et al., 2012; Schauer et al., 2005; Yelle et al., 1991).

To exploit this natural variation, two large and highly polymorphic *S. pennellii* BIL populations were developed using the self-compatible *S. pennellii* accession LA5240 (‘LOST’) as donor and the cultivated *S. lycopersicum* lines LEA (determinate) and TOP (indeterminate) as recurrent parents (Torgeman & Zamir, 2023). Comprising approximately 1400 and 500 lines (Supplemental Figure 1), respectively, these populations offer unprecedented genetic resolution for mapping complex traits while avoiding confounding hybrid incompatibilities associated with other *S. pennellii* accessions.

Here, we used these populations to dissect the genetic architecture of tomato fruit primary metabolism using targeted GC–MS profiling across independent experiments. High-resolution QTL mapping identified loci associated with variation in metabolite abundance, including sugar-related traits. Functional validation was performed using overexpression lines of the invertase inhibitor *SlINVINH3,* which showed allele-specific effects on sucrose content, with one overexpression line showing ∼20% and another ∼80% increase relative to control. Importantly, genome-wide epistasis scans showed non-additive interactions influencing sucrose accumulation, highlighting the role of genetic interactions in shaping fruit metabolism.

## Results

### Metabolic profiling of fruits in a tomato BIL population

To investigate the metabolic composition of tomato fruit in the newly developed LEA and TOP BIL populations and to identify metabolic QTL with higher resolution, we performed comprehensive GC-MS metabolic profiling of red ripening fruits from 1400 LEA and 500 TOP BILs across multiple experiments (Figure 1, Supplemental Table 1). In addition, a further experiment was conducted on both fruit and leaf materials on 50 selected LEA BILs (Supplemental Table 1). These lines represent the metabolic diversity of the LEA population, allowing cross-validation of the large-scale profiling data and estimation of the heritability of metabolic traits.

**Figure 1:**
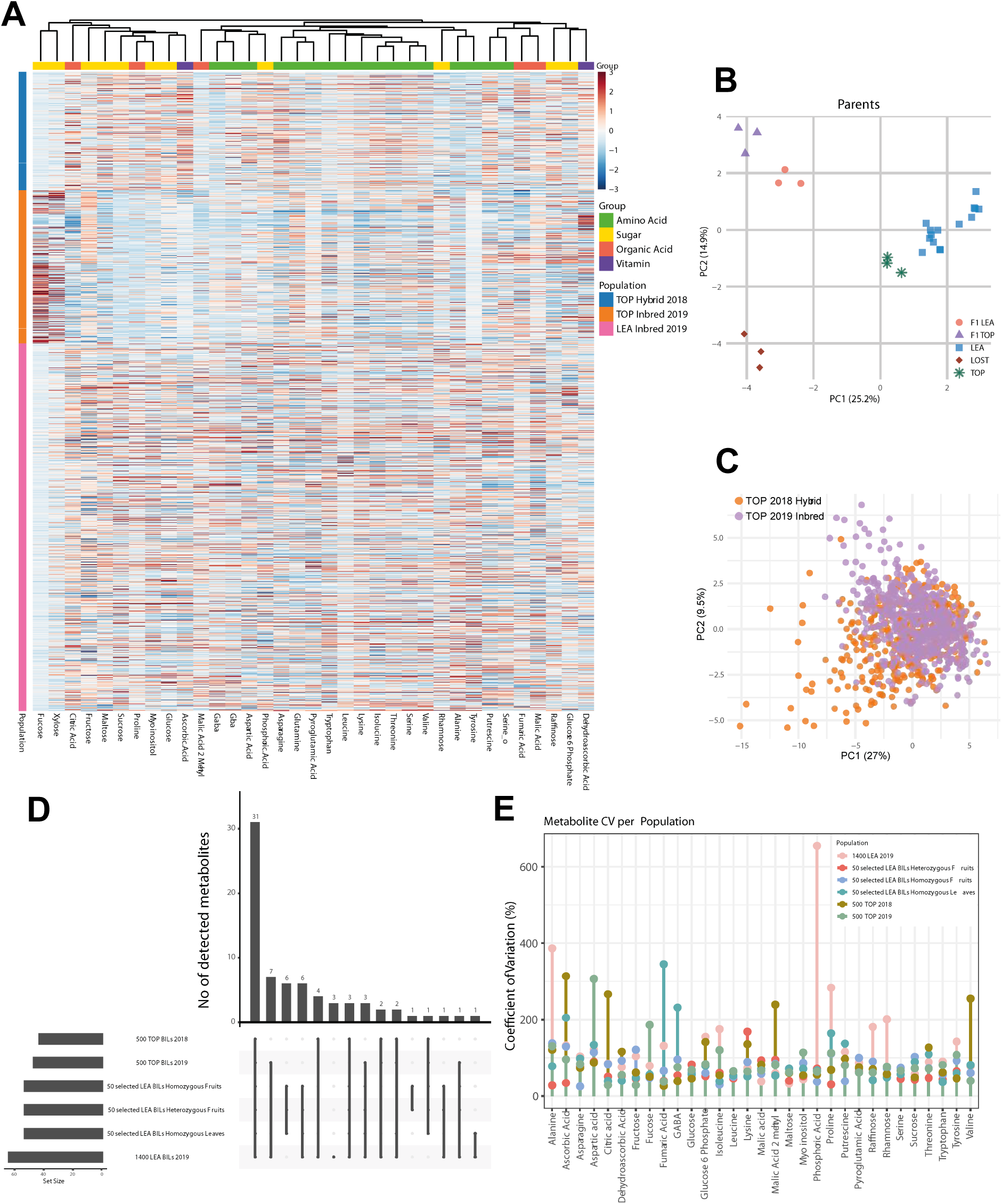
Large-scale metabolite profiling shows shared and population-specific variation in LEA and TOP BIL populations. **(A)** Heatmap showing 35 metabolites’ abundance in TOP and LEA BILs. Internal standard normalized data were then log2-transformed, and subsequently z-score normalized across samples for each metabolite. Hierarchical clustering was applied to metabolites, while samples were ordered by year. Colors represent relative abundance (red=higher, blue=lower). Metabolites are annotated by biochemical class, including sugars, amino acids, sugars, organic acids and vitamins as indicated above the heatmap. **(B)** Principal component analysis (PCA) of primary metabolite profiles illustrates the separation along PC1 and PC2 among parental lines (LEA, TOP, and LOST) and F1 hybrids. LEA is shown as blue squares, TOP as green stars, LOST as brown diamonds, F1 TOP as purple triangles, and F1 LEA as pink circles. **(C)** 500 TOP BILs 2018 and 500 TOP BILs 2019 primary metabolites are illustrated along PC1 and PC2. Data were internally standard-normalized, log10 transformed, and z-score-scaled by dividing each metabolite by its standard deviation prior to visualization. **(D)** UpSet plot showing shared and unique primary metabolites across six populations (1400 LEA BILs 2019; 500 TOP BILs 2018 and 2019; 50 selected homozygous and heterozygous fruits; and homozygous leaves). A core set of 31 metabolites was shared across populations, whereas each population contained only a small number of unique or partially shared metabolites (1-7 per population), indicating limited population-specific metabolite presence. **(E)** Coefficient of variation (CV) analysis of 31 common primary metabolites across 1400 LEA BILs, 500 TOP BILs and 50 selected LEA BILs. CV (%) was calculated across all metabolites and within each population to assess variability.

Before analyzing the full LEA (1400 lines) and TOP (500 lines) BIL datasets, we examined metabolite profiles of the parental lines (LEA, TOP, and the wild species LOST) and their F1 hybrids. A heatmap displaying the relative abundance of 35 shared metabolites across the 1400-line LEA BIL population and two sets of the 500-line TOP BIL population (2018 hybrid and 2019 inbred experiments) was generated, with hierarchical clustering applied to metabolites (Figure 1A). The heatmap revealed distinct patterns of sugar accumulation between populations and years. For example, fucose and xylose levels were elevated in the TOP BILs in 2019 but showed the opposite trend in the TOP BILs in 2018 and in the LEA BILs. Conversely, sucrose accumulation was reduced in TOP BILs in 2019 relative to TOP BILs in 2018 and LEA BILs, where higher levels were observed. Additionally, the amino acids proline, lysine and alanine were lower in the TOP BILs and higher in the LEA BILs.

Principal component analysis (PCA) explained 25.2% (PC1) and 14.9% (PC2) of the variation in the metabolite dataset of the parents (Figure 1B). LEA and TOP were partially separated along PC1, whereas the wild parent (LOST) was clearly separated from both cultivated backgrounds. PCA of the two TOP experiments (Figure 1C) further illustrated this variation: although many lines overlapped, a subset displayed broader dispersion across principal component space, indicating quantitative metabolic divergence among genotypes. PCA of the 50 selected LEA BILs grown in the 2020 field trial, including both inbred (PC1 25.6% and PC2 12.2%) and hybrid (PC1 32.8% and PC2 11.5%) fruits as well as primary metabolites in leaves (PC1 29.1% and PC2 7.6%), is shown in Supplemental Figure 2. The higher PC1 contribution in hybrid fruits suggests that heterozygosity increases metabolic diversity, while the distinct variance patterns in leaves indicate tissue specific regulation of primary metabolism.

### Variation in primary metabolite content across both LEA and TOP populations

To dissect the genetic basis of variation in primary metabolites abundance, we analyzed skinned pericarp from 1400-line LEA BILs harvested in 2019, 500-line TOP BILs harvested in 2018 and 2019 and 50 selected LEA BILs grown in 2020. For the 50 selected LEA lines, both homozygous and heterozygous fruits, as well as the leaves that were collected from the homozygous plants, were sampled and analysed.

Metabolite profiling was performed using GC-MS. In total, we quantified 66 metabolites in the 1400-line LEA dataset and 53 in both fruits and leaves of the 50 selected LEA BILs. For the TOP population, 45 and 47 metabolites were quantified in the 2018 and 2019 harvests, respectively. The detected compounds included sugars and sugar alcohols, organic acids, amino acids and ascorbic acid (vitamin C). Broad variation in metabolite abundance was observed across both the LEA and TOP populations (Figure1).

UpSet analysis revealed a substantial core metabolite set shared across 1400 LEA BILs, 500 TOP BILs and 50 selected LEA BILs, comprising 31 primary metabolites (Figure 1D). In contrast, each population harboured only a small number of unique or partially shared metabolites (ranging from 1 to 7), indicating limited population-specific metabolite presence. This pattern suggests that primary metabolism is largely conserved across populations, with metabolic diversification occurring through a restricted subset of metabolites rather than widespread metabolic reconfiguration. To assess the variability of primary metabolites across 1400 LEA BILs, 500 TOP BILs and 50 selected LEA BILs, we calculated the coefficient of variation (CV, %) for thirty-one common metabolites across all samples and within each population (Figure 1E) indicated that most metabolites exhibit moderate variability (CV 20-150 %) across populations, while certain metabolites display extreme variability in specific populations. For instance, phosphoric acid indicates particularly high CV in the 1400 LEA 2019 population, suggesting population-specific metabolic divergence. Overall, these results show both class-specific and population-specific patterns of metabolite variability.

In the LEA population, metabolite variance ranged from 0.01 to 554.72, substantially higher than previously reported for *S. pennellii* introgression lines (Schauer et al., 2005). In TOP BILs, variance ranged from 0.02 to 149.07 in 2019 and from 0.0015 to 128.92 in 2018. The higher variance in BILs reflects their design: each line carries multiple (∼12) introgressions from wild donor LA5240, compared with a single ILs, resulting in greater recombination and phenotypic diversity. Across populations, organic acids such as malic acid (in LEA and inbred TOP) and ascorbic acid (in hybrid TOP) were among the most abundant metabolites. Sugars (e.g., sucrose, fructose, myo-inositol) and amino acids (e.g., leucine, glutamine, proline) were also highly abundant. Conversely, quinic acid, benzoic acid, and certain sugars (fucose, sucrose, fructose) showed low abundance in both LEA and TOP lines (Supplemental Table 2).

Next, we estimated the broad-sense heritability (*H^2^*) of each metabolite based on fruit from the 50 selected LEA inbred lines (Supplemental Figure 3A). Heritability values ranged from 15% for glucose 5 phosphate to 80% for trehalose. Citric acid (78%) and threonic acid (79%) also showed high variability, whereas malic acid (46%) and ascorbate (65%) displayed moderate heritability. These results indicate that genetic variation contributes substantially to metabolite abundance in tomato fruits and that many metabolites are moderately to highly heritable in this population. The dataset generated here thus provides a valuable resource for further studies on plant metabolism and biosynthetic regulation. In *Arabidopsis*, broad- and narrow-sense heritabilities coincided for 30 primary metabolites, indicating predominantly additive genetic variance, with only 2-oxoglutaric acid showing more than 10% non-additive variance (Rodriguez Cubillos et al., 2018). In wheat, metabolite heritabilities were generally lower (mean ∼25%), although a subset exceeded 40% (Matros et al., 2017). In contrast, large-scale rice metabolomic surveys reported higher values, with 59-68% of metabolites exhibiting heritability above 50%, likely reflecting broader genetic diversity (Chen et al., 2014).

In parallel, SNP-level modes of inheritance were determined by estimating marker effects across 50 selected LEA with replicated inbred and hybrid lines. Allelic effects were classified as additive, dominant, recessive, or overdominant according to the direction and magnitude of the wild allele effect (Supplemental Figure 3B). Across metabolites, inheritance patterns were diverse, with a prominent contribution of non-additive effects. Together, these findings show that primary metabolism in tomato fruit is under substantial genetic control but is shaped by a complex architecture in which dominance and overdominance play major roles alongside additive effects.

### QTL identification for tomato metabolic variation using both LEA and TOP populations

To identify QTL associated with metabolite variation in both populations, a high-density bin map was constructed using almost 8000 SNP markers, in which defined genomic regions from the wild accession *S. pennellii* LA5240 (LOST) replace the corresponding intervals of the cultivated tomato *S. lycopersicum* LEA and TOP backgrounds (Torgeman & Zamir, 2023). These genomic maps, combined with metabolite data, enabled high-resolution QTL mapping.

Using a marker regression approach with Bonferroni-corrected significance thresholds, we detected significant QTL for 55 of 66, 14 of 47, and 19 of 45 metabolites across the genome in the 1400-line LEA BIL, 500-line TOP 2018, and 500-line TOP 2019 populations, respectively (Figure 2A). QTL intervals were defined based on introgression boundaries, encompassing the most significant SNP and adjacent markers marking the transition between *S. pennellii* and *S. lycopersicum* alleles (Figure 2A). Representative genes located within the QTL intervals are indicated in Figure 2A.

**Figure 2:**
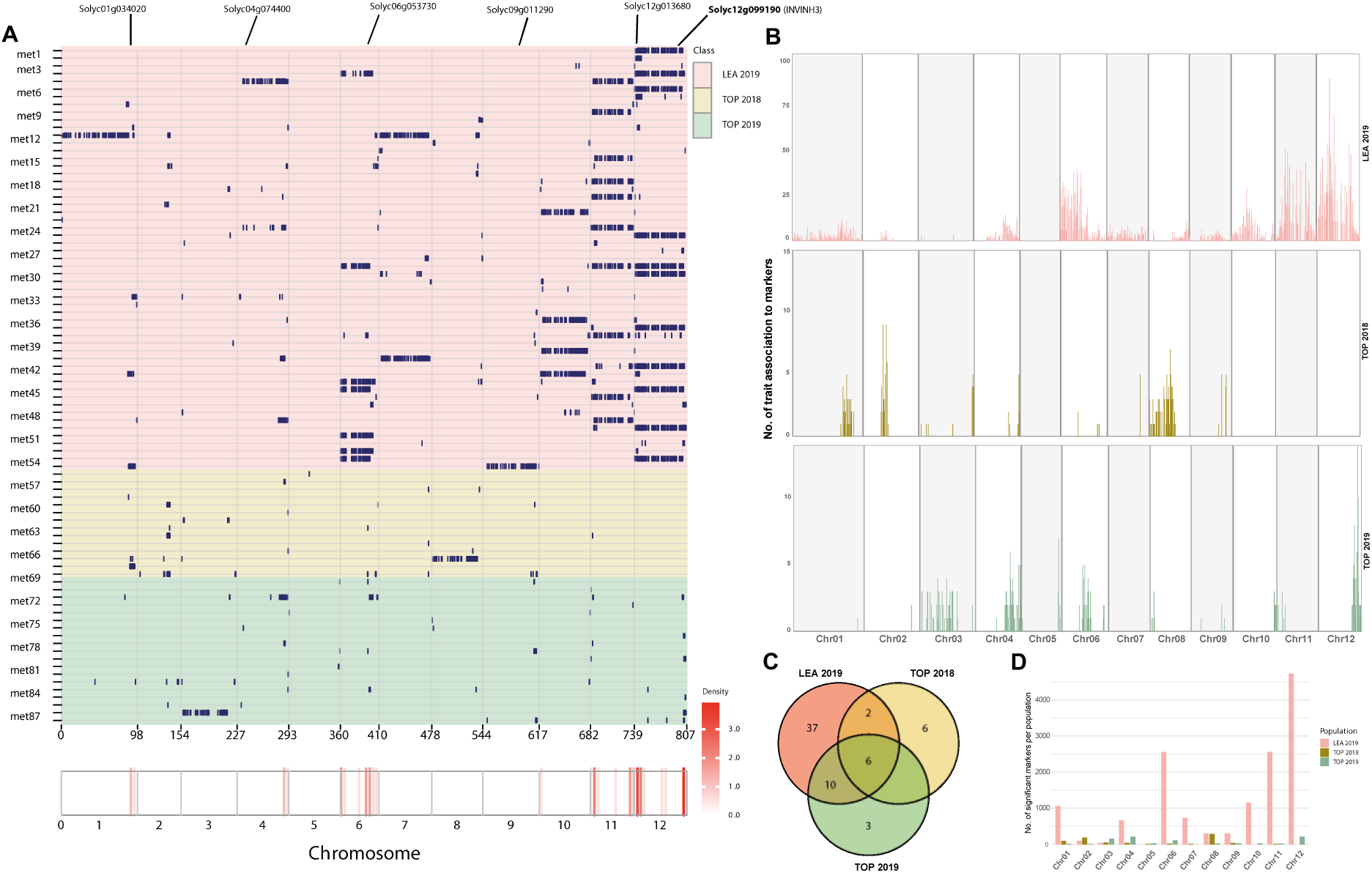
Comparative landscape of primary metabolite QTL across LEA and TOP BIL populations. **(A)** Chromosomal distribution of 55, 14, 19 metabolic QTL identified in the LEA 2019, TOP 2018 and 2019, respectively. QTL across the tomato genome associated with primary metabolite changes from the LEA and TOP populations are shown as dark blue. Each row represents an individual primary metabolite with its significantly associated SNPs. The x-axis indicates the genetic positions across the tomato genome in cM. The threshold of –log_10_(p) was set to 5, representing a Bonferroni corrected statistical threshold. **(B)** Number of traits associated with significant markers for QTL with each primary metabolite class, chromosome-wise for 1400 LEA BILs 2019, 500 TOP 2018 and 500 TOP 2019, respectively. **(C)** Distribution of significant QTLs across three populations (LEA 2019, TOP 2018 and TOP 2019). The Venn diagram shows the number of significant QTLs shared among or unique to each population. Overlapping areas indicate QTL detected in multiple populations, while non-overlapping sections represent population-specific numbers of QTL. This highlights both common and unique genetic loci contributing to trait variation across populations. **(D)** The bar plot indicates the total number of significant traits per population (pink LEA 2019; dark yellow TOP 2018; and green TOP 2019) across all chromosomes.

In the 500-line TOP BILs (2018 and 2019) and the 50 selected LEA BILs (both inbred and hybrid), clear differences in the number and distribution of mQTL were observed. In LEA BIL fruits, a major hotspot on chromosome 12 was detected (Figure 2A), consistent with results in TOP inbred fruits (2019) and the 50 selected LEA inbred and hybrid fruits (Supplemental Figure 4). However, this chromosome 12 hotspot was absent in the leaf metabolic profiles of the 50 selected LEA inbred lines. Chromosomes 6, 11 and 12 contained the highest number of QTL in the full 1400-line LEA population, whereas TOP hybrid fruits (2018) showed the greatest number of QTL on chromosome 2. TOP inbred fruits (2019) again displayed a strong hotspot on chromosome 12 (Figure 2B). The larger 1400-line LEA BIL population showed a greater number of significant marker–trait associations compared with the TOP BIL populations (Figure 2B and 2D). Significant QTL were either shared among populations or unique to individual populations, with population-specific QTL counts and overlapping regions summarized in Figure 2C.

Candidate genes underlying detected mQTL were primarily identified based on high-resolution mapping in the 1400-line LEA population and subsequently cross-checked against the TOP BIL inbred and hybrid datasets (Supplemental Table 3). Most QTL identified here represent previously unreported loci. To assess mapping resolution and accuracy, we compared our results with published *S. pennellii* BIL QTL (Ofner et al., 2016). Several overlaps were observed; for example, QTL for fructose and glucose on chromosome 6 matched previously reported loci, consistent with earlier studies (Schauer et al., 2008; Schauer et al., 2006). These overlaps demonstrate that metabolic variation can be linked to defined genomic loci, in some cases pinpointing genes with known physiological roles, such as *SELF-PRUNING* (Pnueli et al., 1998). A further illustrative case is the H locus, harboring 32 annotated genes, among which Solyc10g078970 is proposed as the putative causal gene (Torgeman et al., 2024).

Our analysis also revealed novel mQTL. For instance, a strong QTL for aspartic acid was detected on chromosome 4 in the LEA population and in two TOP BIL datasets. Multiple candidate genes reside within this interval (Figure 3), suggesting a conserved genetic basis for aspartate metabolism across populations. The interval, defined between markers SSL2.50CH04_60743982 and SSL2.50CH04_61004473, spans approximately 21 annotated genes, including Solyc04g075000, which encodes serine/threonine protein kinase (Supplemental Table 3). The recurrence of this locus across populations suggests a conserved genetic determinant underlying aspartate metabolism. Myo-inositol likewise mapped a shared region on chromosome 4 in the LEA and TOP populations. Raffinose was associated with a common significant SNP on chromosome 6 in TOP 2019 and LEA 2019. Dehydroascorbic acid was detected on chromosome 7 in LEA 2019 and TOP 2018. Finally, xylose was mapped to chromosome 9 in LEA 2019 and TOP 2019.

**Figure 3:**
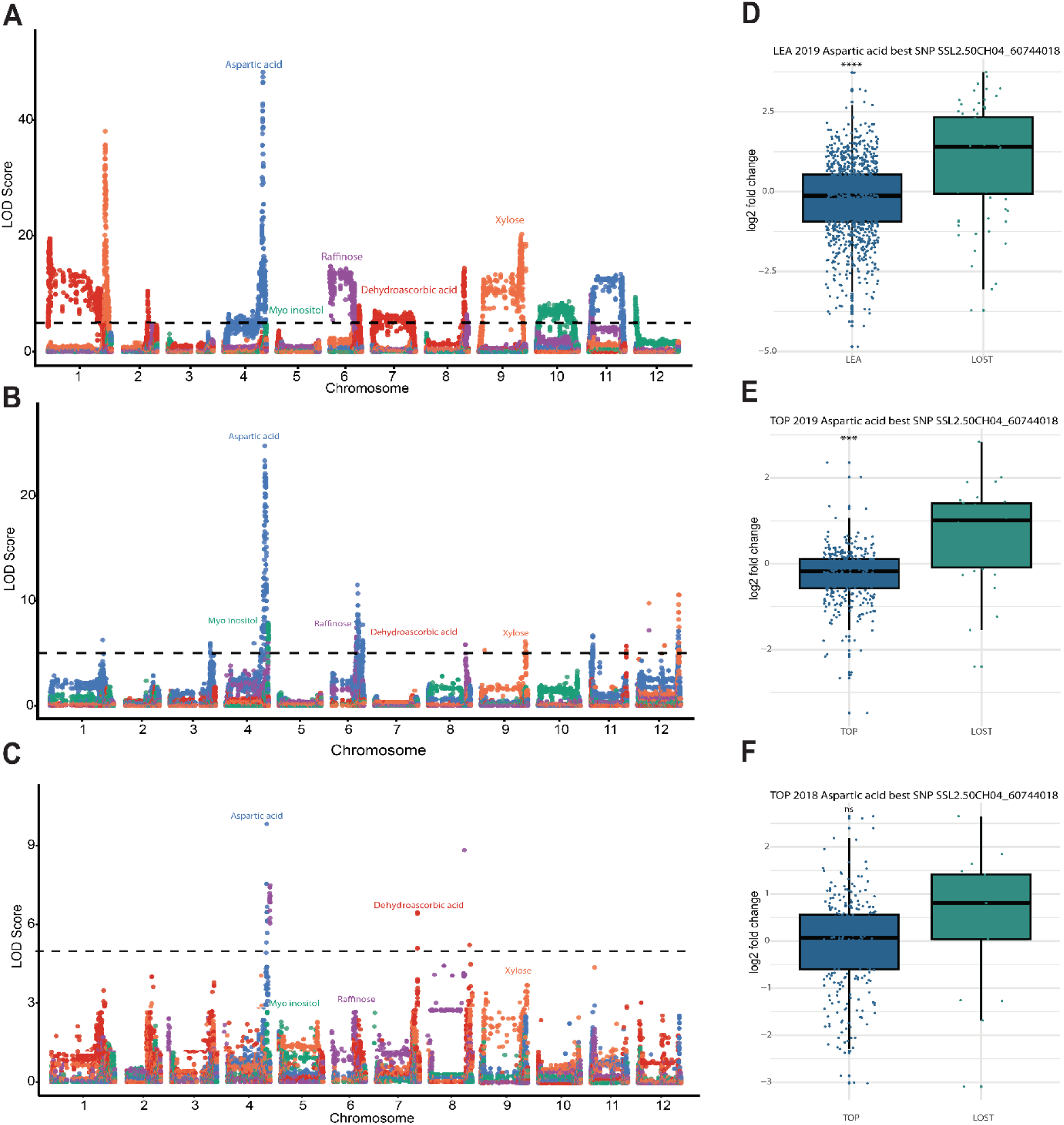
Selected mQTL, including a novel locus associated with aspartic acid accumulation in LEA and TOP BIL populations. **(A, B, C)** Manhattan plots of metabolite QTL results for LEA 2019, TOP 2019 and TOP 2018, respectively. A significant SNP on chromosome 4 (SSL2.50CH04_60744018) is consistently associated with levels in all three datasets. In addition to aspartic acid, other metabolites are displayed in Manhattan plots. Shared significant SNPs were detected for myo-inositol (green) on chromosome 4, raffinose (purple) on chromosome 6, dehydroascorbic acid (red) on chromosome 7 and xylose on chromosome 9. **(D, E, F)** The boxplots display the haplotype distribution of the best SNP (SSL2.50CH04_60744018) for LEA 2019, TOP 2019 and TOP 2018, respectively. The x-axis of each boxplot represents the haplotypes of domesticated alleles as LEA and TOP, as well as wild allele LOST (*S. pennellii* LA5240). The y-axis represents the log_2_ fold change. p value * < 0.05; ** < 0.01; ***< 0.001.

Finally, we assessed correlations between Brix and the levels of hexose sugars and sucrose for 1400-line LEA BILs. Although Brix is primarily influenced by soluble sugars, correlations were weakly positive. Notably, the major QTL for Brix colocalized with sugar-related mQTL on chromosome 6 (Supplemental Figure 6), consistent with previous reports associating this locus with the *SELF-PRUNING* (SP) gene (Torgeman et al., 2024). Brix was further evaluated in greenhouse experiments conducted in 2017, 2018 (both hybrid and inbred) and 2023. In all four datasets, the *S.pennellii* (LA5240) allele at the chromosome 12 QTL peak SNP (SSL2.50_CH1266307289) significantly increased Brix by 7.5%, 3.2%, 4.3% and 8.4%, respectively. Box plots showing the distribution of Brix values by SNP allele are represented in Supplemental Figure 6.

### Mapping of novel mQTL controlling sucrose levels in tomato fruit

A novel QTL for sucrose content was identified on chromosome 12 in the LEA BIL population (Figure 4A), with the lead SNP at SSL2.50CH12_66307289. The QTL peak mapped to a narrow interval between 66.29 Mb and 66.32 Mb, as defined by the boundaries of the wild introgression segment (Figure 4B). Closer inspection of this region revealed multiple annotated genes underlying the QTL peak (Figure. 4C). Haplotype analysis showed that lines carrying the *S. pennellii* allele at this locus exhibited significantly higher sucrose levels than those carrying the *S. lycopersicum* allele (Figure 4 D, E and F), indicating that the wild allele promotes sucrose accumulation.

**Figure 4:**
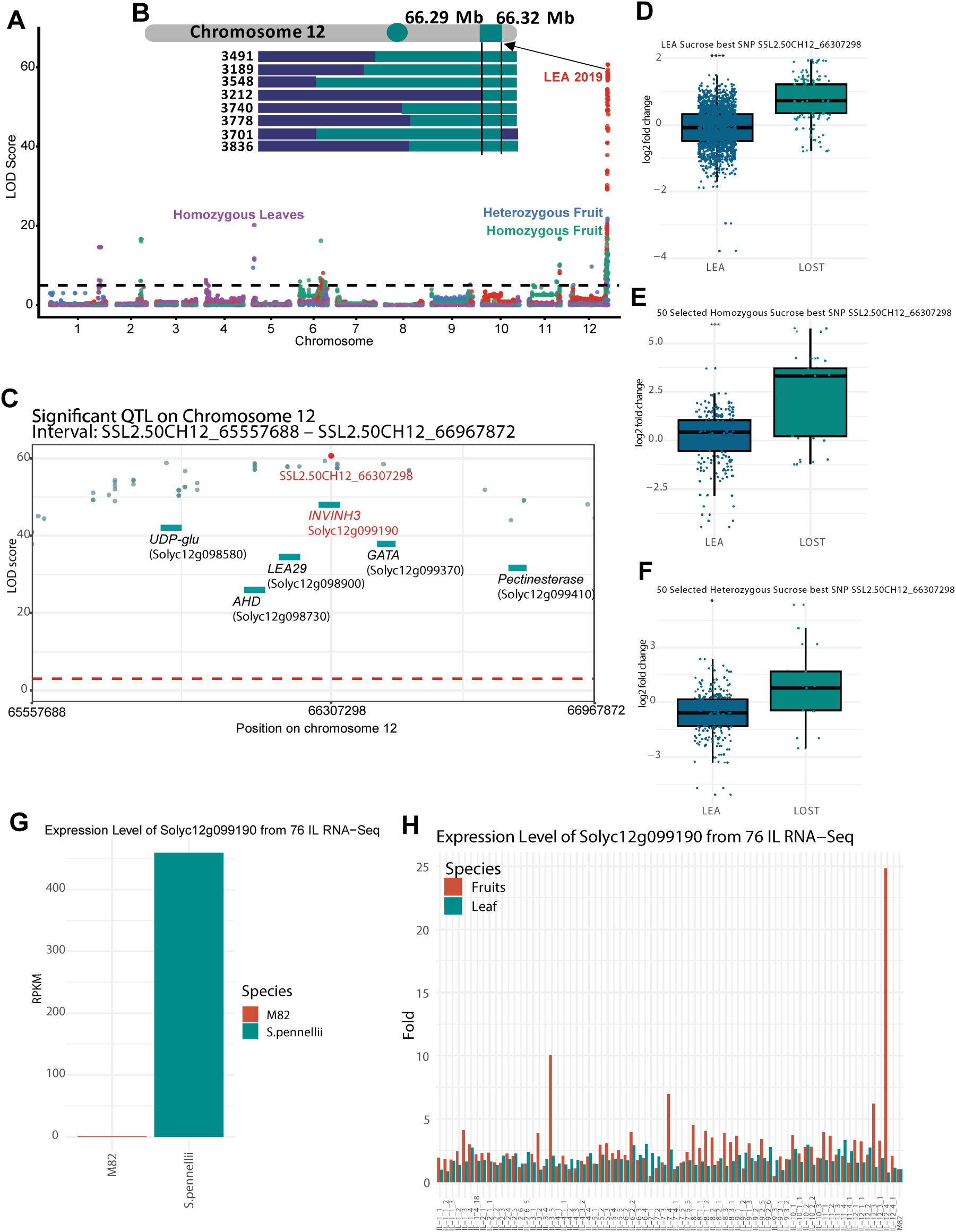
Large-scale mapping reveals a novel sucrose QTL on chromosome 12 in LEA BILs. **(A)** The Manhattan plot indicates the novel QTL on chromosome 12 for sucrose. The best SNP is SSL2.50CH12_66307298 from GC-MS analysis of 1400 BILs (red) and 50 selected fruit inbred (green) and hybrid (blue), however, 50 selected inbred leaf (purple) of LEA BILs did not show a significant QTL on chromosome 12 but rather on chromosome 5. **(B)** The QTL interval in which the invertase inhibitor gene is located, between 66.29 Mb and 66.32 Mb in the region, was observed after checking the starting and ending points of the wild introgressed. **(C)** The major QTL region on chromosome 12. The QTL interval spans from SL2.50CH12_65557688 to SL2.50CH12_66967872. The major associated marker (SSL2.50CH12_66307298) is highlighted in red. Green dots represent other significant SNPs exceeding the significance threshold, which is indicated by the horizontal red line. Genes located within and around the QTL region are shown as green boxes. **(D, E, F),** Boxplots display the haplotype distribution of the significant SNP (SSL2.50CH12_66307298) for 1400 LEA BILs and 50 selected inbred and hybrid fruits, respectively. The x-axis of each boxplot represents the haplotypes of domesticated alleles as LEA and TOP, as well as wild allele LOST (*S. pennellii* LA5240). The y-axis represents the log_2_ fold change. **(G)** The bar plot indicates the expression level of *SlINVINH3* from RNA-seq data of 76 introgression lines (IL) for both fruits and leaves; the data were normalized to M82. **(H)** The expression reads per million mapped reads (RPKM) of *SlINVINH3* for M82 and *S. pennellii* (LA0716) for fruits and leaves. The threshold of –log_10_(p) was set to 5, representing a Bonferroni corrected statistical threshold. p-value * < 0.05, ** < 0.01, *** < 0.001.

This sucrose QTL was also detected in the fruit of the 50 selected LEA inbred and hybrid BILs, confirming the result from the full LEA dataset (Figure 4A). However, no significant association was observed in leaf tissue, indicating that this QTL is fruit-specific. The same sucrose QTL interval was also identified in the TOP inbred BILs (2019), but not in the TOP hybrid BILs (2018) (Supplemental Figure 5A-B).

To determine whether this QTL had been previously reported, we examined several exotic germplasm tomato populations, including, 76 ILs (Schauer et al., 2008; Schauer et al., 2006), 107 *S. neorickii* BILs both homozygous and heterozygous (Brog et al., 2019), 446 *S. pennellii* BILs (Ofner et al., 2016), previously published GWAS datasets (Tieman et al., 2017; Torgeman et al., 2024; Zhu et al., 2018) an unpublished metabolite dataset and 150 accessions from the Bulgarian GWAS tomato diversity panel (unpublished) (Supplemental Figure 5 C, D, E and F). None of these populations displayed a significant QTL for sucrose on the identified loci on chromosome 12.

Together, these results indicate that the sucrose QTL on chromosome 12 is a novel QTL and unique to the LOST *S.pennellii* (5240) wild accession, the fruit-specific locus that is present in both LEA and TOP genetic backgrounds, but absent from previously characterized introgression and GWAS populations.

### Validation of *SlINVINH* candidate genes behind the QTL for sucrose

Following QTL mapping in the 1400-line LEA population, we examined the confidence intervals of significant QTL to prioritize candidate genes for functional validation. A novel QTL on chromosome 12 (SL2.50 position 66291403–66438505) spans ∼147 kb and contains 23 annotated genes, including three coding invertase inhibitor proteins (Solyc12g099190, Solyc12g099200 and Solyc12g099210).

Expression profiles of these invertase inhibitor (*SlINVINH*) genes were examined using fruit and leaf RNA-seq data from *S. pennellii* LA07i6 (Bolger et al., 2014) and 76 introgression lines. All three genes were expressed in *S. pennellii*, but *Solyc12g099190* (*SlINVINH3*) showed markedly higher expression (∼500 RPKM) in fruit and was not expressed in cultivated *S. lycopersicum* cv. M82 (Figure 5B). Similarly, RNA-seq data from the introgression lines indicate that *SlINVINH3* expression is approximately 25-fold higher in IL 12-4 compared to other ILs and M82 (Figure 4H).

**Figure 5:**
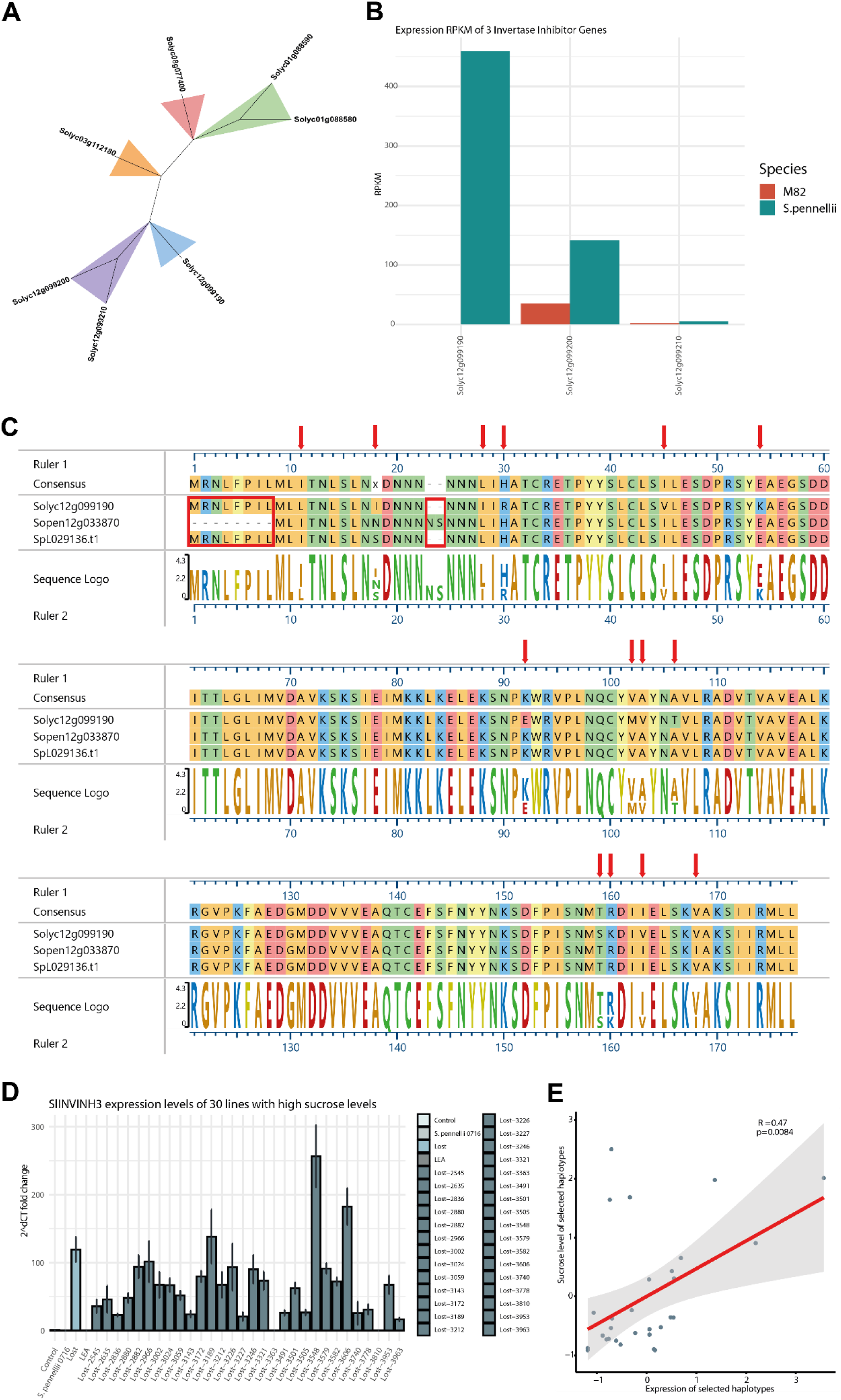
Integrated phylogenetic and expression evidence implicates *SlINVINH3* in sucrose accumulation. **(A)** In the phylogenetic tree of invertase inhibitor genes, the candidate gene is highlighted in blue on the tree. **(B)** The fruit expression data from RNA-seq of 76 ILs of three homologous invertase inhibitor genes located on chromosome 12, indicated that Solyc12g099190 is highly expressed in *S. pennellii* (LA0716) compared to the other two of the invertase inhibitor genes. **(C)** The protein sequence comparison of Solyc12g099190 with its alleles Sopen12g033870 (*S.pennellii* LA0716) and SpL029136.t1 (LOST) indicates that there are polymorphisms and deletions between domesticated and the other two *S.pennellii* alleles. **(D)** Expression levels of invertase inhibitor of the 30 wild-type haplotypes with the highest sucrose levels were checked with qRT-PCR. Additionally, the expression level of invertase inhibitor was checked in the parental lines LEA, as domesticated and LOST (*S.pennellii* 5240) and *S.pennellii* LA0716 as wild-type parents. The comparison was done with the Moneymaker as a control plant. **(E)** The correlation analysis was performed between the qRT-PCR results of selected haplotypes and their sucrose levels.

To further assess tissue specificity, we examined TomExpress, which showed higher expression of *SlINVINH3* in *S. lycopersicum* roots relative to fruit, whereas Tomato eFP browser (Fernandez-Pozo et al., 2017) indicated strong expression of *SlINVINH3* in mature fruit of *S. pennellii* (LA0716), while the two-cell wall invertase inhibitors (Solyc12g099200 and Solyc12g099210) showed negligible fruit expression (Supplemental Table 4). Based on its strong fruit-specific expression and position within the sucrose QTL, *SlINVINH3* (Solyc12g099190) was prioritized as the most likely causal gene.

Phylogenetic analysis of invertase inhibitor protein sequences revealed seven invertase inhibitor loci in tomato (Figure 5A). Sequence comparison of the *SlINVINH3* coding regions revealed two potential ATG start codons in *S. lycopersicum* and S.pennellii (LA5240), with translation predicted to initiate from the first ATG. In contrast, in *S. pennellii* (LA0716), the annotated coding sequence initiated from the second ATG, resulting in an 18 bp shorter CDS corresponding to a six amino acid truncation at the N-terminus (Figure 5C). Notably, the upstream ATG in LA0716 is located within the 5’ untranslated region, suggesting allele-specific differences in translation initiation. The signal peptide was checked for *SLINVINH3* for three alleles using SignalP 6.0 (Teufel et al., 2022). The results indicated that signal peptide (Secretory/Signal peptidase I) was 0.5318, 0.4458 and 0 for *S. lycopersicum* and *S. pennellii* (LA5240) and *S. pennellii* (LA0176), respectively (Supplemental Table 5).

The cis-regulatory elements of *SlINVINH3* were analyzed using 1 kb upstream promoter sequences from *S. pennellii* (LA0716), *S. lycopersicum* and *S. pennellii* (LA5240) via New PLACE (Higo et al., 1999). The sucrose-responsive motif SREATMSD motif (TTATCC) (Zolotarov & Strömvik, 2015), was present in both *S.pennellii* (LA0716) and *S. pennellii* (LA5240) promoters but absent in *S. lycopersicum*. WRKY transcription factor binding motifs, including WBBOXPCWRKY1 (TTTGACY), WBOXNTERF3 (TGACY) and WRKY71OS (TGAC) were identified in all promoters. GATABOX (GATA) motifs were present in all three promoters at distinct positions. Tandem repeats analysis using PlantPAN 4.0 (Chow et al., 2024) indicated three repeat elements (ATA, AATAATAATAATAACAAC, AATAATAATAACAACAACAAC) in both *S.pennellii* alleles (LA0716 and LA5240), whereas these repeats were absent in *S. lycopersicum* (Supplemental Table 5).

To test this hypothesis, stable overexpression lines were generated carrying three alleles: the domesticated LEA allele and two wild-type alleles from *S. pennellii* LA0716 and LA5240. Effects on fruit sucrose accumulation and ripening were evaluated.

In parallel, 30 LEA-derived BILs carrying LOST haplotypes associated with high sucrose were analyzed. qRT-PCR in fruit pericarp revealed that 28 of 30 lines exhibited high *SlINVINH3* expression relative to parental LEA and LOST lines, *S. pennellii* LA0716, and Moneymaker controls (Figure 5D). Expression positively correlated with sucrose content (r = 0.47) (Figure 5E). Finally, qRT-PCR of T1 OX lines confirmed significantly elevated *SlINVINH3* expression in both leaves and fruit in all LOST-allele OX lines compared to the LEA or *S. pennellii* alleles (Supplemental Figure 7).

### *SlINVINH3* is a key gene influencing sugar metabolism in tomato fruit

To further elucidate the role of *SlINVINH3* in sugar metabolism and to biologically validate our mapping results, we generated *SlINVINH3* overexpression lines driven by the cauliflower mosaic virus 35S promoter (*SlINVINH3*-OX). Three *SlINVINH3* alleles were overexpressed, including the domesticated LEA allele and two wild alleles from *S. pennellii* accessions LA0716 and 5240 (Lost). Following Agrobacterium-mediated transformation and molecular confirmation, one independent LEA-OX line, two Lost-OX lines and two Pen-OX lines were selected for subsequent phenotypic and metabolic analyses.

At early stages, all OX lines flowered earlier than wild-type controls and most also initiated fruit set earlier (Supplemental Figure 8). To evaluate broader developmental effects, progeny from T1 transformants were phenotyped throughout growth under greenhouse conditions. Traits recorded included flower number, open flowers, fruit number and shoot biomass (Supplemental Figure 9).

OX lines produced smaller fruits on average. A reduction in mean fruit weight was observed in Lost OX lines, whereas Pen OX lines showed a modest increase. However, neither average nor total fruit weight differed significantly from wild-type plants (Supplemental Figure 9A-B). Measurements of Brix total soluble solids (TSS) revealed a decrease in Lost OX lines and a relative increase in Pen OX lines, although these differences did not reach statistical significance (Supplemental Figure 9C).

Because the primary goal of this study was to elucidate the genetic control of primary metabolites in tomato fruit, metabolic profiling focused on this tissue. A total of 56 primary metabolites were quantified in fruit and leaf tissues, with 50 metabolites shared between both tissues. These shared metabolites were used for hierarchical clustering and partial least squares discriminant analysis (PLS-DA) to assess global metabolic variation.

In fruit, hierarchical clustering revealed a distinct metabolic signature in two Lost OX lines (Lost 17 and Lost 7). Several organic acids formed a prominent cluster and were markedly reduced in these lines, whereas amino acids and sugars showed increased accumulation (Figure 6A). This inverse relationship between organic acids and sugars/amino acids was a dominant feature of the clustering pattern and was not observed in the remaining OX lines or the wild-type control.

**Figure 6:**
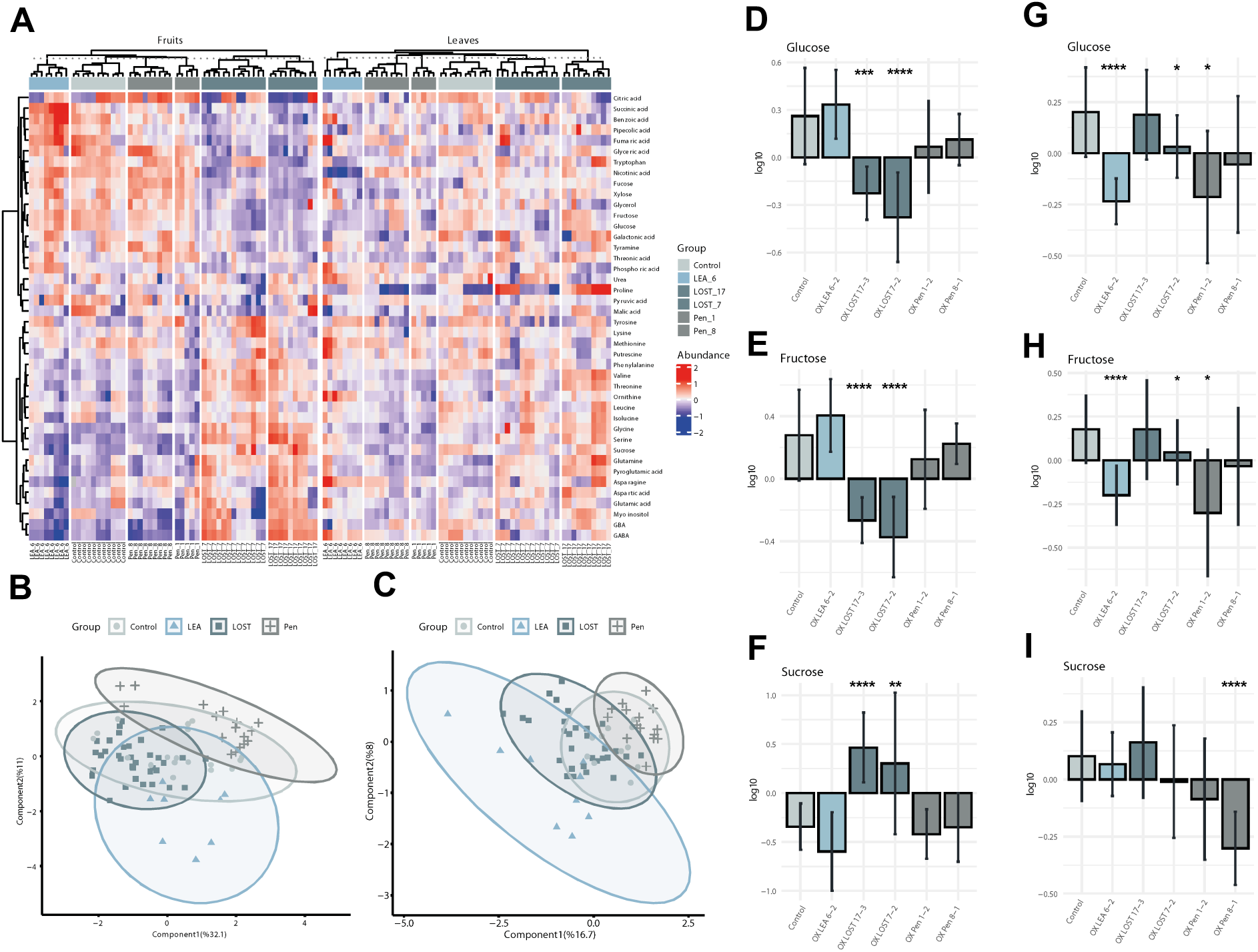
Metabolomic analysis of fruits and leaves from *SlINVINH3* overexpression lines. **(A)** Heatmap illustrates log_10_ fold change of the relative abundance of sugars, amino acids and organic acids in both fruits and leaves. **(B)** Partial least squares discriminant analysis (PLS-DA) of fruit metabolite profiles across all analytical platforms. Component 1 (32.1%) and component 2 (11%) are shown on the x- and y- axis, respectively. **(C)** PLS-DA of leaf metabolite profiles, with component 1 at (16.7%) and component 2 at (8%) shown on the x- and y- axis, respectively. Groups indicate the OX lines of distinct alleles and MM as a control for PLS-DA. **(D, E, F)** Glucose, fructose and sucrose distribution of fruits of OX lines and the control plants. **(G, H, I)** Glucose, fructose and sucrose distribution of leaves of OX lines and the control plants. p value *<0.05, **<0.01, ***<0.001. circle = control, triangle = LEA (as domesticated allele), square = LOST (*S. pennellii* 5240 allele), plus = Pen (*S. pennellii* 0716). 2-3 biological replicates of each line and 3 technical replicates of each biological replicate were used to generate the data. Outliers were removed while generating the figures.

PLS-DA of fruit metabolites showed that the first two components explained 32.1% and 11.0% of the total variance, respectively (Figure 6B). In leaves, the first two components accounted for 16.7% and 8.0% of the variance (Figure 6C). Despite detectable metabolic shifts, no clear genotype-based separation was observed in either tissue, indicating that the effects of *SlINVINH3* overexpression were metabolite-specific rather than globally discriminative.

Analysis of soluble carbohydrates revealed pronounced allele-specific effects. In ripe fruit, sucrose levels were significantly increased in the Lost OX lines (Lost 17 and Lost 7), whereas LEA and Pen OX lines did not differ from wild-type controls (Figure 6F). Conversely, glucose and fructose levels were significantly reduced in the same Lost lines, with no consistent changes detected in other OX lines (Figure 6D-E).

In leaf tissue, sucrose levels were significantly reduced only in the Pen 8 line. Glucose and fructose levels were decreased in LEA 6, Lost 7, and Pen 1, while the remaining lines showed no significant differences relative to wild type (Figure 6G-I).

Together, these results validate *SlINVINH3* as a key gene underlying the chromosome 12 sucrose QTL. Notably, enhanced sucrose accumulation in ripe fruit was observed exclusively in Lost OX lines, indicating that coding sequence polymorphisms present in the Lost allele are likely functionally important for sucrose regulation. Although the specific causal polymorphisms remain to be identified, these findings suggest that allelic variation in *SlINVINH3* contributes to natural variation in fruit sucrose content. We note that differences in transgene expression levels may also contribute to phenotypic outcomes, as transcript abundance in Pen OX lines was significantly lower than in Lost OX lines.

### Epistasis

To demonstrate the value of the newly developed populations and to uncover hidden non-additive interactions between QTLs, we dissected the role of epistasis in shaping primary metabolism. Epistatic interactions were identified using a two-dimensional QTL scan testing all pairwise combinations of loci, enabling the detection and quantification of interacting QTLs affecting metabolite accumulation. Epistasis emerged as a pervasive feature of the genetic architecture in all populations, encompassing both more than additive and less than additive effects (Supplemental Table 6).

The LEA BIL population exhibited a greater number of epistatic interactions than the TOP populations, likely reflecting its larger size (1400 lines and 500 lines), which increases statistical power to detect interactions and captures a wider range of allelic combinations. Additional contributions may arise from differences in genetic diversity, population structure, and environmental conditions. Epistasis analyses were restricted to digenic combinations represented by at least ten individuals carrying both wild-species alleles, to mitigate bias from segregation distortion. Across all populations, we identified 208 epistatic interactions, of which 125 were less-than-additive and 83 were more-than-additive, with most showing modest effect sizes.

Comparison across growing seasons in the TOP population revealed limited stability of interactions, indicating a strong environmental influence. In 2019, 20 metabolites exhibited epistasis, with GABA showing the highest interaction frequency, while most metabolites had a single interaction. In 2018, only 10 metabolites exhibited interactions, with ascorbic acid, sucrose, tyrosine, and calystegine A showing the most interactions. Only a small subset, including sucrose, raffinose, maltose, glucose-6-phosphate, and xylose, displayed epistasis in both years (Figure 7A). In the LEA population, epistatic interactions were detected for 53 metabolites, with alanine, calystegine A, glyceric acid, and malic acid-2-methyl exhibiting the highest number of interactions (Figure 7B). Digenic epistatic interactions were further visualized on a circos physical map (Figure 7C).

**Figure 7:**
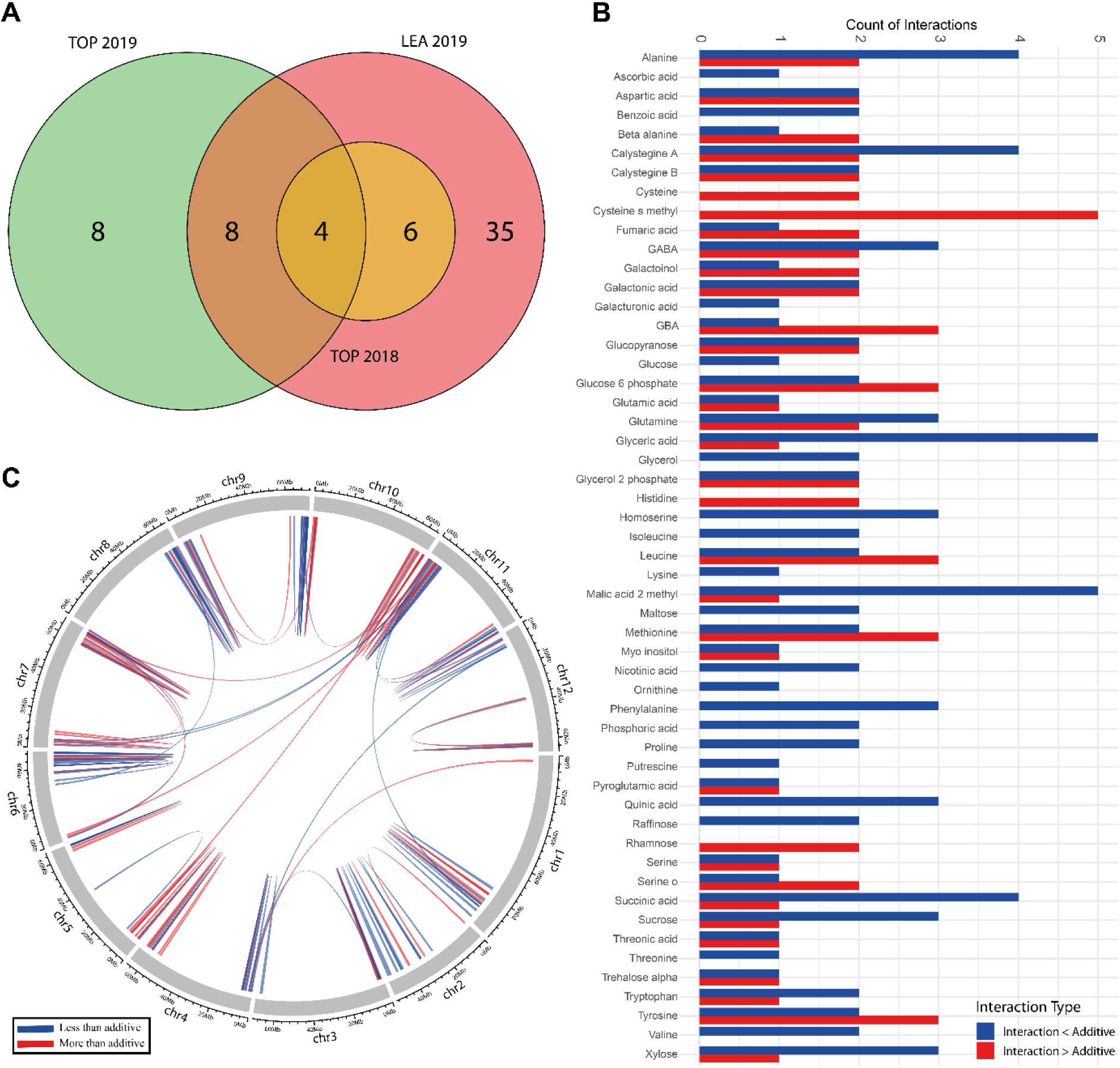
Population-dependent epistasis among metabolic QTL. **(A)** Venn diagram represents the number of metabolites that were detected commonly and individually for epistatic interactions for 500-line TOP 2018, 500-line TOP 2019 and 1400-line LEA 2019 BIL populations. **(B)** The bar plots represent the 53 primary metabolic traits, which indicate epistatic interactions for 1400-line LEA BILs. The blue color represents the less-than-additive interactions, the red color represents more-than-additive interactions. **(C)** Circos plot indicates the chromosomal positions of epistatic interactions of 1400-line LEA 2019 for 53 primary metabolites. Significant epistasis was defined using an LOD_int_>3, indicating strong evidence for interacting QTL. We used LOD int>3 (Broman & Sen, 2009) to indicate significant epistasis.

Several genomic regions functioned as epistatic hotspots. For example, for LEA BIL population on chromosome 6 (SSL2.50CH06_41586962), multiple metabolites, including GABA, malic acid-2-methyl, and phenylalanine, showed interactions. Interestingly, the direction of epistasis varied: GABA displayed more-than-additive effects, whereas malic acid-2-methyl and phenylalanine were less-than-additive, suggesting metabolite-specific modulation by the same locus. Candidate genes within this interval included transcription factors from the MYB (Solyc06g066180), WRKY (Solyc06g066370), ERF (Solyc06g066540), and bHLH (Solyc06g066580) families, supporting a central regulatory role.

Specific loci in the TOP population further illustrate epistatic control. A shared interaction region on chromosome 7 (SSL2.50CH07_66509915) in TOP 2019 affected glucose-6-phosphate and maltose. This interval contains a glucose-6-phosphate/phosphate translocator (Solyc07g064270), a key mediator of plastid-to-cytosol carbon exchange. Glucose-6-phosphate exhibited more-than-additive interactions, while maltose showed less-than-additive effects, highlighting metabolite-specific epistatic modulation. In TOP 2018, chromosome 10 (SSL2.50CH10_61307233) harbored interactions for tyrosine and xylose, with F-box (Solyc10g080020) and MADS-box (Solyc10g080030) proteins as candidate genes. Tyrosine showed more-than-additive and xylose less-than-additive effects, indicating a regulatory role mediated via protein degradation and transcriptional control.

Collectively, these results demonstrate that environmental variation strongly shapes epistatic architecture, and although individual epistatic effects are generally smaller than major additive QTLs, their cumulative impact is likely substantial, particularly for metabolites with multiple interacting loci. This underscores the importance of non-additive genetic variance in determining primary metabolite variation.

## Discussion

In this study, the genetic basis of natural variation in tomato fruit primary metabolism was investigated using two *S. pennellii* (LA5240) BIL populations, LEA and TOP, comprising approximately 1400 and 500 lines, respectively (Torgeman et al., 2024; Torgeman & Zamir, 2023). Compared with the previously described *S. pennellii* (LA0716) BIL panel of 446 lines (Ofner et al., 2016), these populations provide substantially increased recombination density and mapping resolution, enabling a more comprehensive dissection of metabolic trait architecture.

Targeted GC–MS profiling of polar metabolites detected 66 metabolites in the LEA population and 45–47 metabolites across two independent harvests of the TOP population. Integration of metabolite data from 50 selected LEA BILs provided an in-depth view of fruit primary metabolism. Metabolite heritability spanned a broad range and was classified as high (> 0.6), moderate (0.4-0.6) or low (< 0.2). While these estimates are based on a single year with three replicates and are therefore expected to vary across environments and seasons (Schauer et al., 2008; Schauer et al., 2006) clear trends were observed. In agreement with earlier studies, metabolites belonging to related biochemical pathways often exhibited comparable heritability, suggesting shared genetic control of pathway-level regulation. In addition to heritability estimates, SNP-level analyses indicated diverse modes of inheritance across metabolites, with a substantial contribution of non-additive effects, including dominance and overdominance. This indicates that, alongside additive genetic variance, non-additive interactions play an important role in shaping primary metabolism in tomato fruit.

The exploitation of wild tomato diversity through introgression and backcross populations has been instrumental in uncovering the genetic architecture of complex traits. In tomato, ILs derived from diverse wild relatives have enabled the systematic mapping of loci underlying fruit quality, yield and metabolic traits (Alseekh et al., 2013; Rousseaux et al., 2005; Schauer et al., 2008; Schauer et al., 2006; Stevens et al., 2008). The *S. pennellii* IL population developed in the M82 background has been particularly influential, leading to the cloning of landmark loci such as *fw2.2* for fruit weight and *Brix9-2-5* for sugar yield, the latter corresponding to allelic variation in the cell wall invertase gene *LIN5* (Frary et al., 2000; Fridman et al., 2004). These studies established sugar metabolism as a central target of domestication and breeding. Building on this foundation, QTL mapping in the LEA and TOP BIL populations identified multiple loci associated with variation in primary metabolite levels. Among these, a major QTL for aspartic acid on chromosome 4 was linked to several putative candidate genes, providing targets for future functional validation.

Consistent with this framework, QTL mapping in the LEA and TOP BIL populations identified multiple genomic regions associated with variation in primary metabolite levels. Among these, we detected a novel sucrose-associated QTL on chromosome 12 encompassing three invertase inhibitor genes *SlINVINH3*, *INVINH1*, and *INVINH2*. To our knowledge, no QTL has previously been reported for this gene cluster. Recent genome-editing studies targeting *INVINH1* and *INVINH2* demonstrated increased soluble solid content in tomato fruit (Kawaguchi et al., 2021), supporting the functional relevance of invertase inhibitor genes in sugar regulation.

We focused further analyses on *SlINVINH3*, as this gene is expressed in tomato fruit, whereas *INVINH1* and *INVINH2* show primarily leaf-specific expression in the cultivated background. Although expression data for LA5240 are not yet available, the absence of this QTL in earlier *S. pennellii* IL and BIL populations suggests that it is specific to the LA5240 accession. Functional validation through overexpression of three *SlINVINH3* alleles (domesticated, LA5240-Lost and LA0716-Pen) showed allele-specific phenotypic effects. All transgenic lines exhibited significantly elevated *SlINVINH3* transcript levels, with the highest expression observed in Lost allele lines. These lines flowered earlier than controls, consistent with the role of the sucrose signalling molecule in floral induction (Cho et al., 2024). Other vegetative and reproductive traits were largely unaffected, indicating a specific rather than pleiotropic developmental effect.

Corresponding to these shifts in sugar composition, total soluble solids (Brix) were significantly reduced in Lost allele overexpression lines, despite no change in fruit weight or number. Metabolite profiling revealed a marked increase in fruit sucrose accompanied by a decrease in glucose and fructose, consistent with enhanced inhibition of invertase activity. These results align with the established role of invertase inhibitors in regulating sucrose cleavage and hexose accumulation (Greiner et al., 1998; Hothorn et al., 2004; Jin et al., 2009). Previous studies identified *SlINVINH3* as a vacuolar invertase inhibitor with elevated expression during fruit ripening (Qin et al., 2016) and silencing of invertase inhibitors has been shown to increase hexose accumulation in tomato fruit (Jin et al., 2009). Our results extend these findings by demonstrating allele-specific functional variation in *SlINVINH3* that contributes to natural diversity in fruit sugar composition and affects Brix.

A major finding of this study is the influence of the chromosome 12 sucrose-associated QTL on fruit Brix across multiple datasets. In the 1400-line LEA BIL population and greenhouse experiments conducted in 2017, 2018 (hybrid and inbred) and 2023, the *S.pennellii* (LA5240) allele at the QTL peak SNP (SSL2.50_CH1266307289) consistently increased Brix by 3-8%, demonstrating a robust effect on total soluble solids. These population-level results complement the functional analysis of *SlINVINH3*, linking allele-specific sugar modulation to measurable improvements in fruit quality.

From a practical perspective, the Brix effect has important implications for both the fresh-market and processing tomatoes. In fresh-market varieties, sugar composition strongly influences flavor and consumer preference, suggesting that allele-specific manipulation of *SlINVINH3* could improve taste. In processing tomatoes, where soluble solids determine product yield and consistency, the fruit-specific invertase inhibitor allele from LA5240 represents a valuable genetic resource. The observed additive and epistatic interactions further indicate that its phenotypic effect depends on genetic background, which should be considered in breeding programs.

Beyond single-locus effects, our findings underscore the widespread role of epistasis in shaping tomato fruit metabolite variation. Two-dimensional scans revealed extensive non-additive interactions, including both more-than-additive and less-than-additive effects, affecting multiple metabolites and differing across populations and years. For instance, glucose-6-phosphate and maltose in TOP 2019 exhibited opposing epistatic directions at a shared locus on chromosome 7, while tyrosine and xylose in TOP 2018 were similarly regulated by a chromosome 10 locus harboring F-box and MADS-box genes. In the LEA BIL population, a shared locus on chromosome 6 harbored multiple TFs, suggesting a regulatory hotspot. These results indicate that genetic interactions modulate metabolite levels in a metabolite-specific manner, and that epistatic effects can mask or enhance the contribution of individual loci in single-QTL analyses.

In conclusion, this study provides a comprehensive dissection of the genetic architecture underlying tomato fruit primary metabolism. By leveraging the high-resolution LEA and TOP BIL populations, we identified numerous loci controlling metabolite variation, including novel QTLs affecting sugar composition, and validated allele-specific functional effects for key genes. Our analyses further reveal that both additive and non-additive interactions contribute to metabolic diversity, reflecting the complex regulatory networks that coordinate pathway-level flux. These findings emphasize the importance of integrating population-scale mapping, functional genomics, and genetic interaction analyses to understand natural variation. Collectively, they offer valuable insights for crop improvement, highlighting genetic strategies to optimize fruit quality and metabolic composition.

## Methods

### Plant material and growth conditions

The two BIL populations, LEA and TOP, used in this study were recently developed (Torgeman et al., 2024; Torgeman & Zamir, 2023). Supplemental Figure 1 represents the schematic overview of crosses to generate the LEA and TOP BILs. Briefly, the tomato wild species *S. pennellii* and domesticated determinate LEA were crossed and an F1 population was generated. The F1 hybrid was backcrossed to both LEA and TOP as the recurrent parents to obtain BILs; thus, the TOP BILs segregate for three possible alleles. BC1 plants were backcrossed to both LEA and TOP parents and around 2000 BC2 plants were generated. After six times of BC2, 500-line TOP BILs and 1400-line LEA BILs were selected and subjected to genotyping by single primer enrichment technology (SPET) (Barchi et al., 2019).

### Metabolite profiling by GC-MS

Fruit materials of both TOP BIL populations which were grown in the greenhouse trial and LEA BIL populations which were grown in the field trial, as well as the leaf material of 50 selected LEA BILs, were ground using the mixer mill. Aliquots of 50 mg leaf and 120 mg fruit were used for metabolite extraction as described before (Salem et al., 2020). Samples were extracted in 1 ml methyl-*tert*-butyl ether (MTBE)-MeOH that contains 20 *µ*L ribitol (20 mg mL^-1^ water), 125 µL isovitexin (1 mg mL^-1^ water) and 45 µL phosphatidylcholine (PC) (1 mg mL^-1^ water) as quantification standards (Salem et al., 2020). In order to prevent the enzymatic conversion of the metabolites, the buffer was kept cold while performing metabolite extraction. First, 1 ml of extraction buffer was added to the frozen leaf samples and vortexed until it was fully suspended. Then the samples were incubated for 10 minutes at a 4 °C orbital shaker. Afterward, the samples were incubated for another 5 minutes in an ultrasonication bath and 500 µl of the water: methanol (3:1) mixture was added into each tube and mixed well. Next, the tubes were centrifuged at full speed for 5 min in a tabletop centrifuge at 4 °C. The second buffer (water: methanol) caused a phase separation. After performing metabolite extraction, 100 µL from the polar phase was taken and concentrated using the Speed Vac and subjected to gas chromatography-mass spectrometry (GC-MS) analysis.

Metabolite analysis by GC-MS was carried out by a method modified from (Lisec et al., 2006). Briefly, concentrated samples were derivatized for 120 min at 37°C in 55 *µ*L of 30 mg mL^-1^ methoxyamine hydrochloride in pyridine, followed by a 30 min treatment with 110 µL of *N-methyl-N-* (trimethylsilyl) trifluoroacetamide (MSTFA) (20 µl fatty acid methyl esters-FAME) at 37°C. GC was performed using a 30-m MDN-35 capillary column. The injection temperature was set at 230°C. Helium was used as the carrier gas at a 2 mL/min flow rate. Mass spectra were recorded at 20 scans/min with a mass-to-charge ratio of 70 to 600 scanning range.

### Chromatogram processing & annotation & normalization

Peak picking from GC-MS chromatograms of 1400-line LEA BILs and 500-line TOP BILs was performed manually and Xcalibur Version 4.2.47 (ThermoFisher Scientific; Waltham, MA, U.S.A.) was used to calculate peak area. Furthermore, the Golm Metabolome database was utilized to cross-reference the mass spectra (Kopka et al., 2005; Lisec et al., 2006).

Normalization of a raw peak area for 1400-LEA BILs, both of 500-line TOP BILs experiments, 50 selected LEA lines and overexpression lines was performed to the batch median. Extreme outlier samples were excluded from the analysis subsequent to PCA.

### QTL mapping, epistasis and heritability

Single QTL mapping analysis was performed using R/qtl marker regression (Broman & Sen, 2009). The bins were clearly defined and a uniform logarithm of odd (LOD) value was assigned for each bin. The confidence interval for each QTL was assigned as 5-LOD based on the Bonferroni calculation.

For the detection of epistasis, two-dimensional, two-QTL genome scans were performed with the integration of a normal model by considering two-dimensional genome scans in the context of normally distributed traits (Broman & Sen, 2009). In the normal model, it is considered that precisely two QTL exist, and each pair of positions in the genome is the putative location for those QTL.

The null model (H_0_), that there is no QTL, is nested in the single-QTL models. Additionally, the additive model (H_α_), they are assumed to act additively, is nested into the full model in which two QTL are allowed to interact. Therefore, the interaction LOD score was tested by comparing the residual of the full model containing all single-QTL effects and two-locus interaction effects with that of the reduced model containing all single-QTL effects but excluding the two-locus interaction effects.

Broad sense heritability was calculated using the data generated from fruit materials from 50 BILs selected from the LEA BIL population, which were grown in the field trial, each line with three replications and a random-effects ANOVA according to the following Eqn (1);

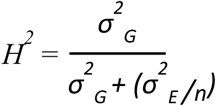

where variance components *σ*^2^_G_, the genetic variance due to the effect of segregant and *σ*^2^_E_, the error variance and *n* is the number of replicates taken for a particular genotype. This was implemented using the “lmer” function in the lme4 R package (Bates et al., 2015).

To determine SNP-level modes of inheritance, marker effects were estimated using phenotypic data from 50 selected LEA BILs for which both inbred and hybrid fruits were available. Genotypic data consisted of SNP markers coded according to genotype classes (1, 2, 3). For each SNP–metabolite combination, individuals with missing values were excluded, and mean metabolite levels were calculated for the three genotype classes.

Additive (a) and dominance (d) effects were calculated based on genotype means, where the additive effect was defined as α =(*μ*_DD_−*μ*_HH_)/2 and the dominance effect as *d*=*μ*_DH_−(*μ*_DD_+*μ*_HH_)/2, with *μ*_DD_, *μ*_DD_, and *μ*_DH_ representing the mean metabolite values of the two homozygous and the heterozygous genotypes, respectively. The ratio *d*/α was then used to classify the mode of inheritance for each SNP.

Inheritance modes were categorized according to the magnitude of the *d*/α ratio: additive (|d/a| < 0.2), partial dominance (0.2–0.8), complete dominance (0.8–1.2), or overdominance (|d/a| > 1.2), with the direction indicating whether dominance favored the domesticated or wild allele. SNPs lacking sufficient genotype representation were excluded from the analysis.

### Generation of overexpressed tomato plants

An overexpression construct for *Sl-INVINH3* (Solyc12g099190) was produced by cloning the respective full-length cDNA pool generated from the fruit pericarp of LEA (domesticated-528 bp), *S. pennellii* 0716 (510 bp) and *S. pennellii* 5240 (528 bp) into pK2GW7 (Karimi et al., 2002) downstream of the 35S promoter subsequent cloning into the binary vector pDONR207. All PCR-based constructs were verified by sequencing. The sequencing-verified vectors were transformed to the competent *A. tumefaciens* GV2260 as previously described (Zhang et al., 2020). Plant transformation was performed by the greenteam of the MPIMP by regeneration of cotyledon segments of *S. lycopersicum* MoneyMaker to calli on a sterile medium, followed by co-cultivation with *A. tumefaciens* GV2260 (Deblaere et al., 1985).

### Growth condition of overexpressed (Zhu et al.) tomato plants

Selection of transformant plants was performed by cutting the upper section of well-rooted plants and placing them on fresh sterile media containing agar (6.8% w/v), MS, sucrose (2% w/v) and 50 *µ*g mL^-1^ Kanamycin. Selection was applied 3 times with 4-6 weeks of cultivation under sterile conditions. Putatively transformed plants were cultivated under sterile conditions using a tissue culture chamber (York International/Johnson Controls; Cork, Ireland) supplying ∼35 µmol m^-2^ s^-1^ of light in a (16h/8h)-(22°C/22°C)-(day/night) cycle. Standard greenhouse conditions were used for the positively selected plants. At the last step of transferring leaf samples were collected for DNA extraction as described before (Lu, 2011). The presence of the transgene was confirmed by PCR with 35S and specific primers, the list of the primers is in Supplemental Table 7.

After harvesting seeds from the T0 plants, the seeds were grown on fresh sterile media containing agar (6.8% w/v), MS, sucrose (2% w/v) and 50 *µ*g mL^-1^ Kanamycin. Since the media contains Kanamycin antibiotics, the negative seeds could not grow and genotyping was performed by PCR with 35S and specific primers with the positive plants.

### Phenotyping of overexpressed tomato plants

Distinct phenotypic traits were recorded during the growth and at harvest time. Flowering time, number of flowers and number of fruits were documented. Red fruits were harvested as well as the total fruit weight was measured by a scale. The average fruit weight per plant was determined by dividing the total fruit weight per plant by the number of fruits. Brix was measured for mature fruits with a digital refractometer. Fresh shoots were dried after harvesting all the fruits for 3 days at 65°C to estimate the shoot dry weight.

### Tissue harvest of overexpressed tomato for metabolic profiling

Young leaf samples were collected before noon (10.00 am to 11.30 am). Per plant, 2 leaflets from young leaves were pooled and treated as one sample. Red ripe fruits were harvested before noon from all plants and the pericarp was taken from one to two fruits per plant on the same day. All samples were frozen in liquid nitrogen and kept at −80°C until the next processing. The fruits were collected from 2-3 biological replicates of each line and 3 technical replicates were used from each biological replicate while performing the data analysis.

### RNA extraction and qRT-PCR

Total RNA was extracted using a Nucleospin RNA isolation kit. One µg of total RNA was used to synthesize the first strand of cDNA. Genomic DNA digestion and cDNA synthesis were performed using the PrimeScript™ RT reagent kit (TaKaRa). Expression of the invertase inhibitor (*SlINVINH3*) genes was analyzed by real-time qRT-PCR using the fluorescent intercalating dye SYBR Green in an iCycler detection system (Bio-Rad; http://www.bio-rad.com/). The specific primers are listed in Supplemental Table 7.

### Heat maps, PCA and PLS-DA

Heat maps were created by clustering the metabolites hierarchically. Principal component analysis (PCA) was performed to visualize the data globally analyze the relationships between the overall traits and determine the traits’ distinctiveness. Partial least squares discriminant analysis (PLS-DA) was performed for the overexpressed lines. log_10_ transformation was applied in addition to mean-centering and divided by the square root of the standard deviation of each variable after internal standard normalized data.

### Phylogenetic Analysis

For phylogenetic analysis, the Phytozome v13 database (Goodstein et al., 2012) was surveyed for *Solanum lycopersicum* invertase inhibitor sequences. Tree reconstruction was performed by the Maximum likelihood algorithm in MEGA 11 (Tamura et al., 2021) with pairwise deletion subsequent to protein sequence alignment. Branches were supported by bootstrapping following 500 replications.

### Computational analysis, packages used and data visualization

R statistical software 4.5.2 was used in the RStudio environment with the help of qtl, ggplot2, wrapr, ggpatern, tidyverse, ggpubr, lme4, dplyr, tibble, complexheatmap, pheatmap and dendsort (Dowle et al., 2019; Kassambara, 2018; Wickham, 2016; Wickham et al., 2019).

## Author contributions

SA conceptualized the work. EK and SA wrote the manuscript with input from all authors. EK performed the experiments and all the data analyses. MWA and ST collected plant materials. DQ processed the LEA-selected population for the fruit. ST and DZ generated the BIL populations and provided the materials as well as the genotypic data. BU provided the sequence data for *S. pennellii*.

## Acknowledgements

SA acknowledges the European Regional Development Fund through the Program “Research Innovation and Digitalisation for Smart Transformation” 2021–2027, Grant No. BG16RFPR002-1.014-0003-C01. And the NATGENCROP project: HORIZON-WIDERA-2022-TALENTS-01, No. 720101087091

## Supplemental Figures

**Supplemental Figure 1:**
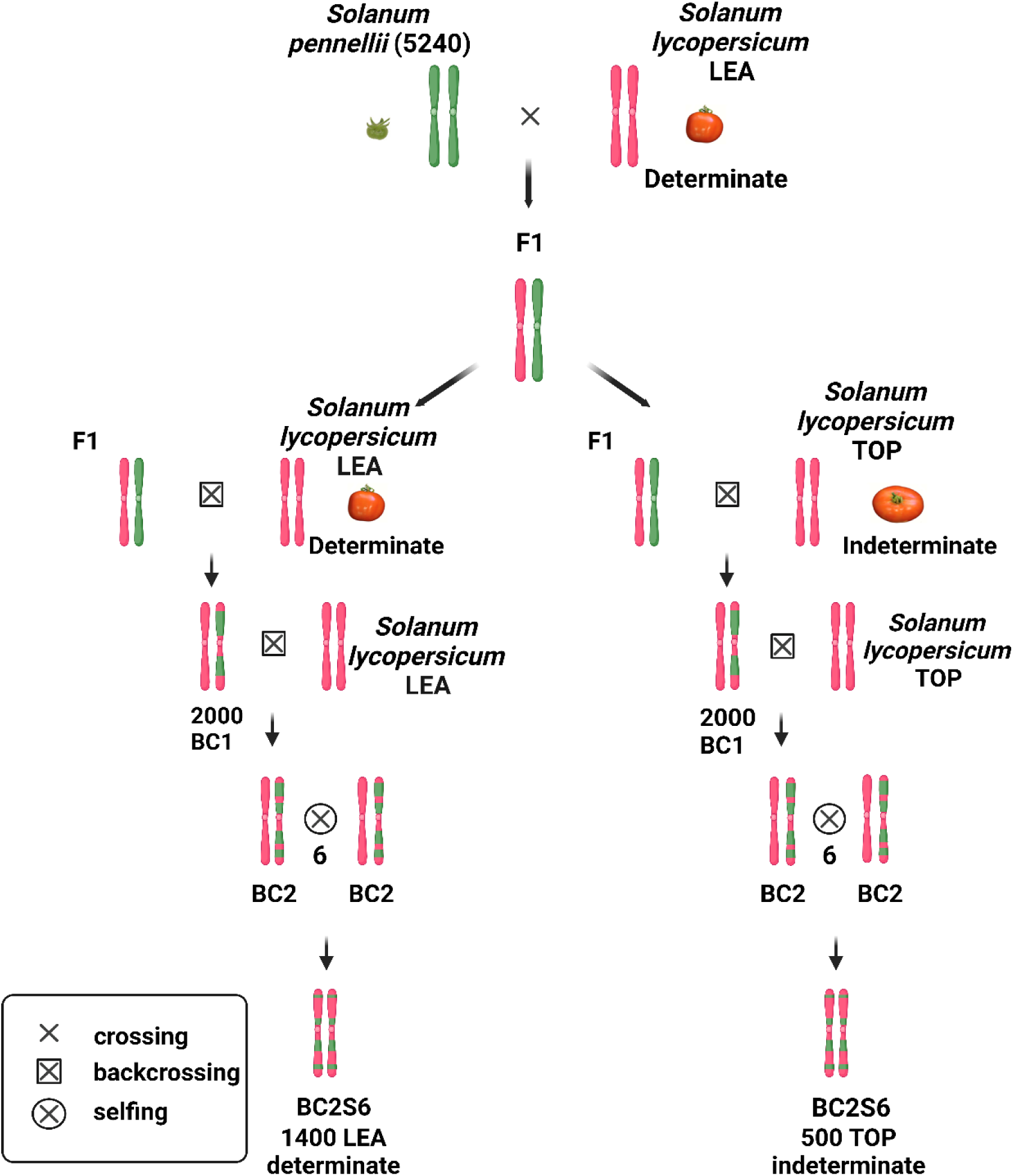
Breeding scheme of both LEA and TOP backcross inbred lines (BILs) populations. Note that a single plant of the hybrid of LEA and LA5240 gave rise to both the LEA and TOP BILs.

**Supplemental Figure 2:**
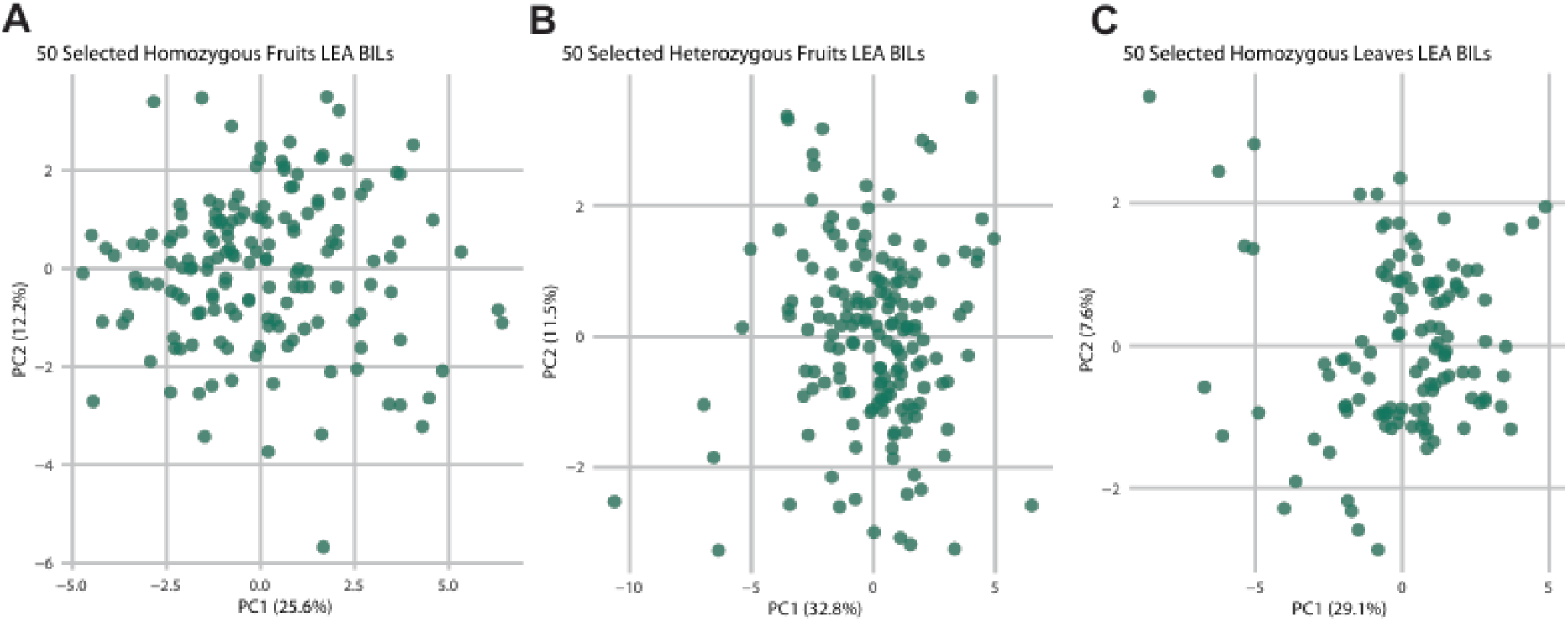
Principal component analysis (PCA) of metabolite profiles from 50 selected LEA BILs. (A, **B)** PCA of 50 selected LEA BILs homozygous and heterozygous for fruit. PC1 and PC2 explain 25.6% and 12.2% of the variance for homozygous fruits, and 32.8% and 11.5% for heterozygous fruits. **(C)** PCA of 50 selected LEA BILs homozygous leaves with PC1 and PC2 explaining 29.1% and 7.6% of the variance. Data were log10-transformed, and scaled by the square root of the standard deviation for each variable after internal standard normalization.

**Supplemental Figure 3:**
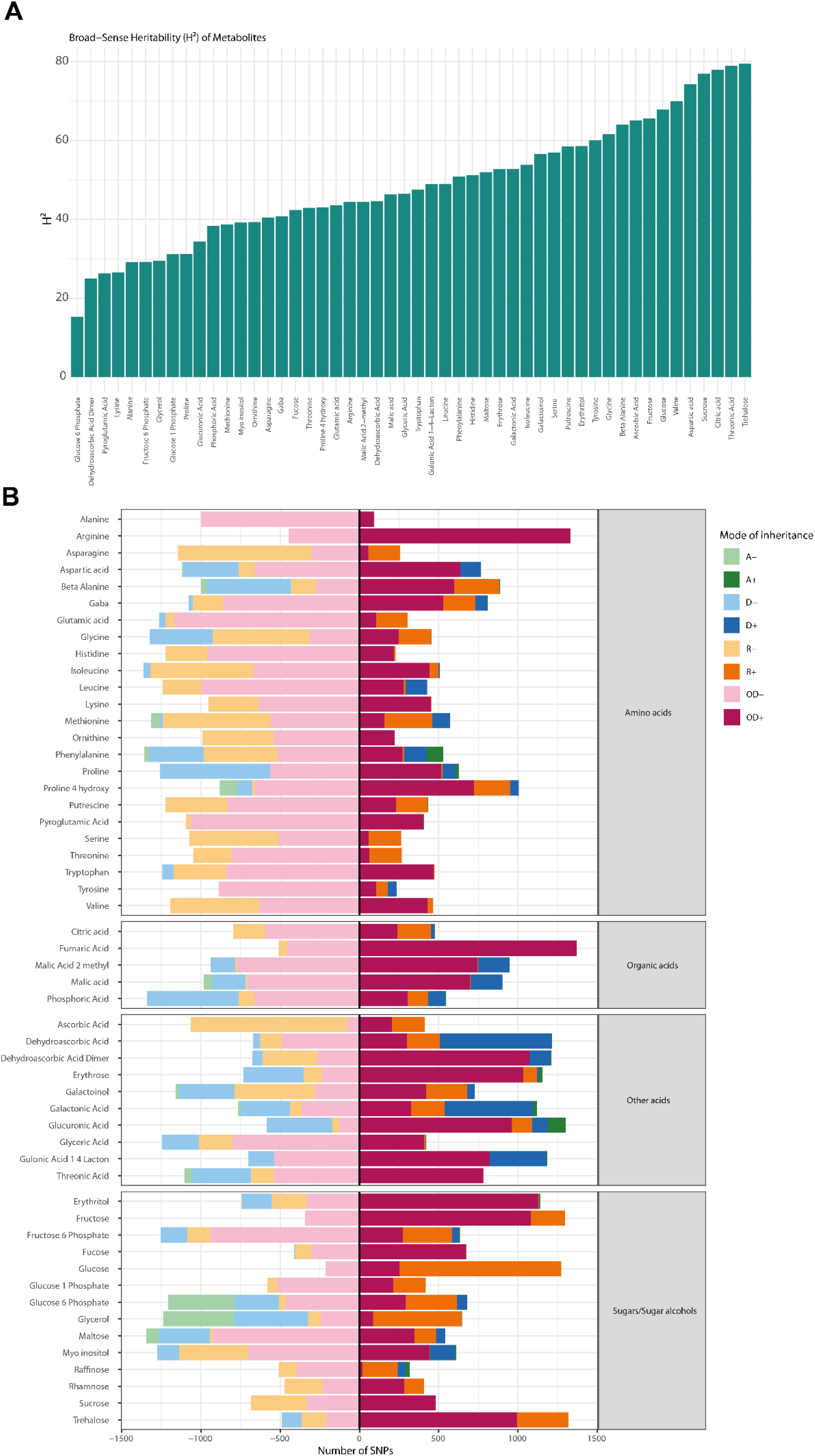
Mode of Inheritance and Broad-Sense Heritability of Metabolites in 50 Selected LEA BIL Population. **(A)** SNP-level modes of inheritance were determined across 27 tomato lines using 50 selected LEA BILs, three replicates per homozygous and heterozygous genotype. Marker effects were estimated at the population level to classify allele action as additive (A+ and A−), dominant (D+ and D−), recessive (R+ and R−), or overdominant (OD+ and OD−). Positive signs (+) indicate that the wild allele increases the trait value, whereas negative signs (−) indicate a decrease. The x-axis shows the number of SNPs assigned to each inheritance category. The vertical line separates decreasing (left) and increasing (right) effects of the wild allele for metabolites. **(B)** Broad-sense heritability (H²) of metabolites was estimated using data from 50 LEA backcross inbred lines (BILs) evaluated in a field trial with three biological replicates per line. Variance components were obtained from a random-effects ANOVA model. Heritability estimates ranged from 15% for glucose 6 phosphate (lowest) to 80% for trehalose (highest), indicating substantial variation in the genetic contribution to metabolite accumulation across traits.

**Supplemental Figure 4:**
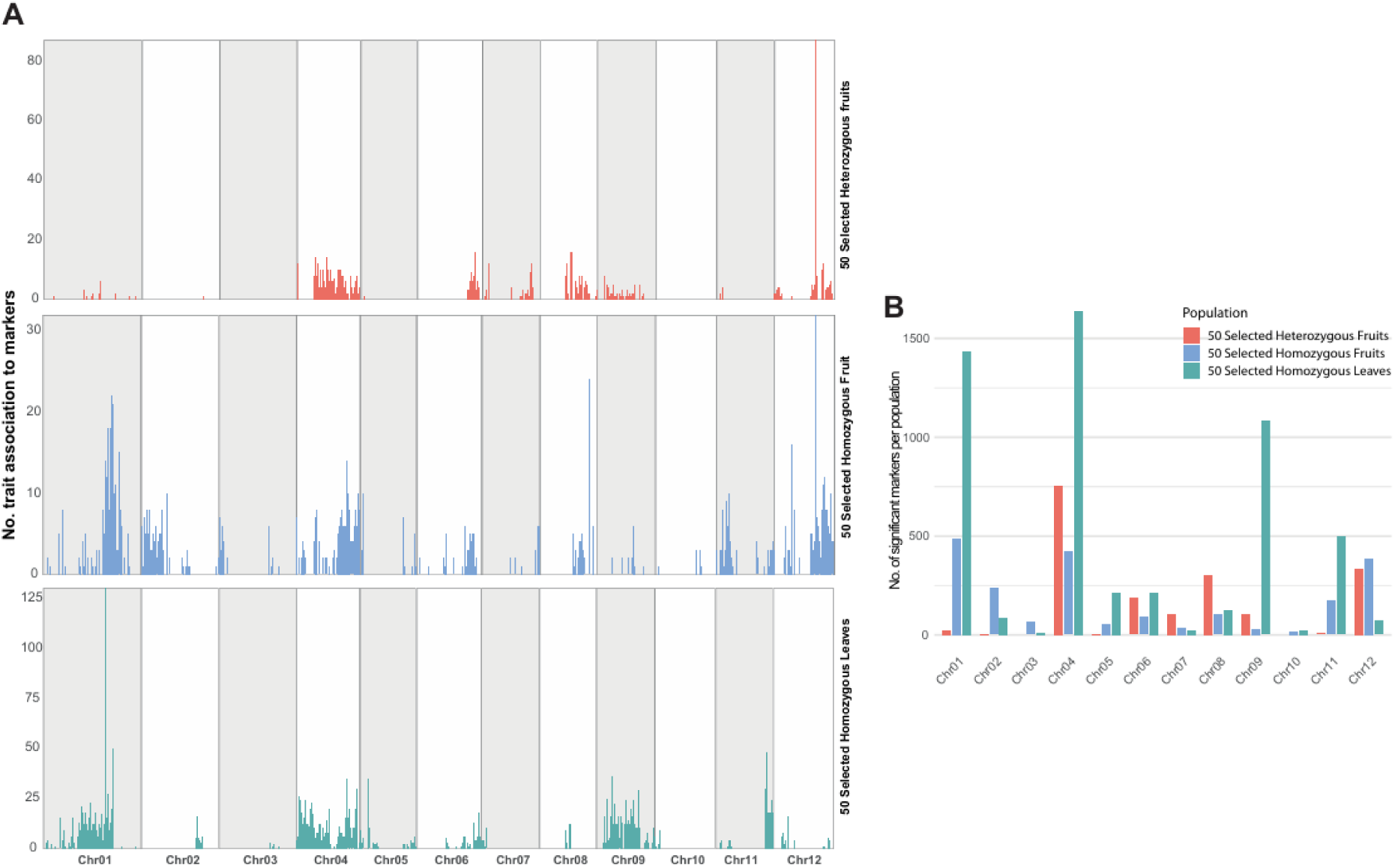
Distribution of QTL-associated markers across chromosomes in the 50 selected LEA BIL populations. (**A)** Bar charts showing the number of traits associated with significant QTL markers on each chromosome in 50 selected LEA BILs, analyzed separately for heterozygous fruits, homozygous fruits, and homozygous leaves. **(B)** Bar plot showing the total number of significant markers per chromosome in the 50 selected LEA BIL population, combining heterozygous and homozygous fruits and homozygous leaves.

**Supplemental Figure 5:**
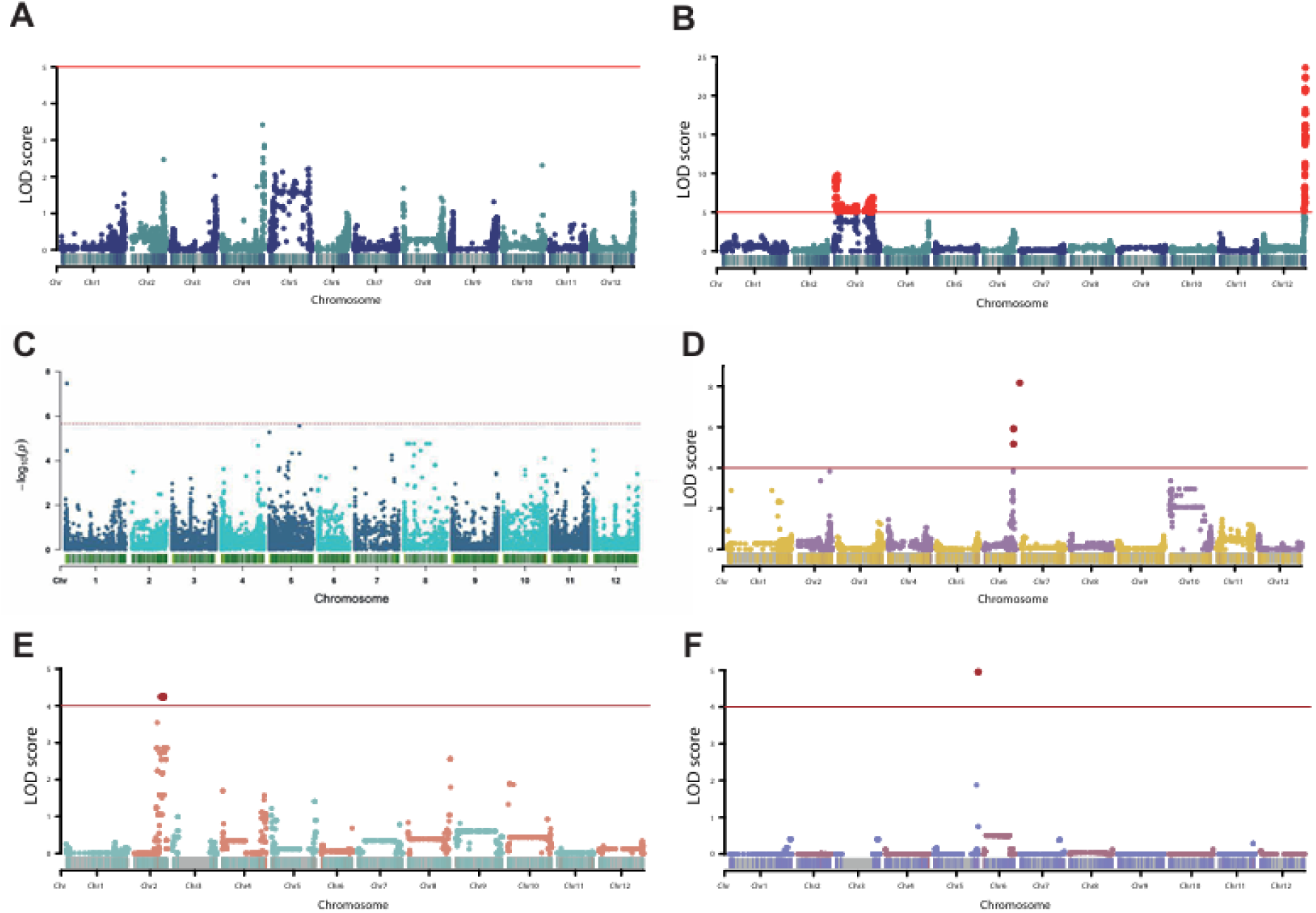
Genome-wide association of sucrose content across tomato populations. **(A)** 500 TOP BILs 2018 (heterozygous). **(B)** 500 TOP BILs 2019 (homozygous) where the significant QTL is located on chromosome 12. **(C)** 150 lines of GWAS Bulgarian population (unpublished). **(D)** *S. pennellii* (0716) 446 BIL population (unpublished). **(E)** *S. neorickii* 107 BILs homozygous. **(F)** *S. neorickii* 107 BILs heterozygous (Brog et al., 2019).

**Supplemental Figure 6:**
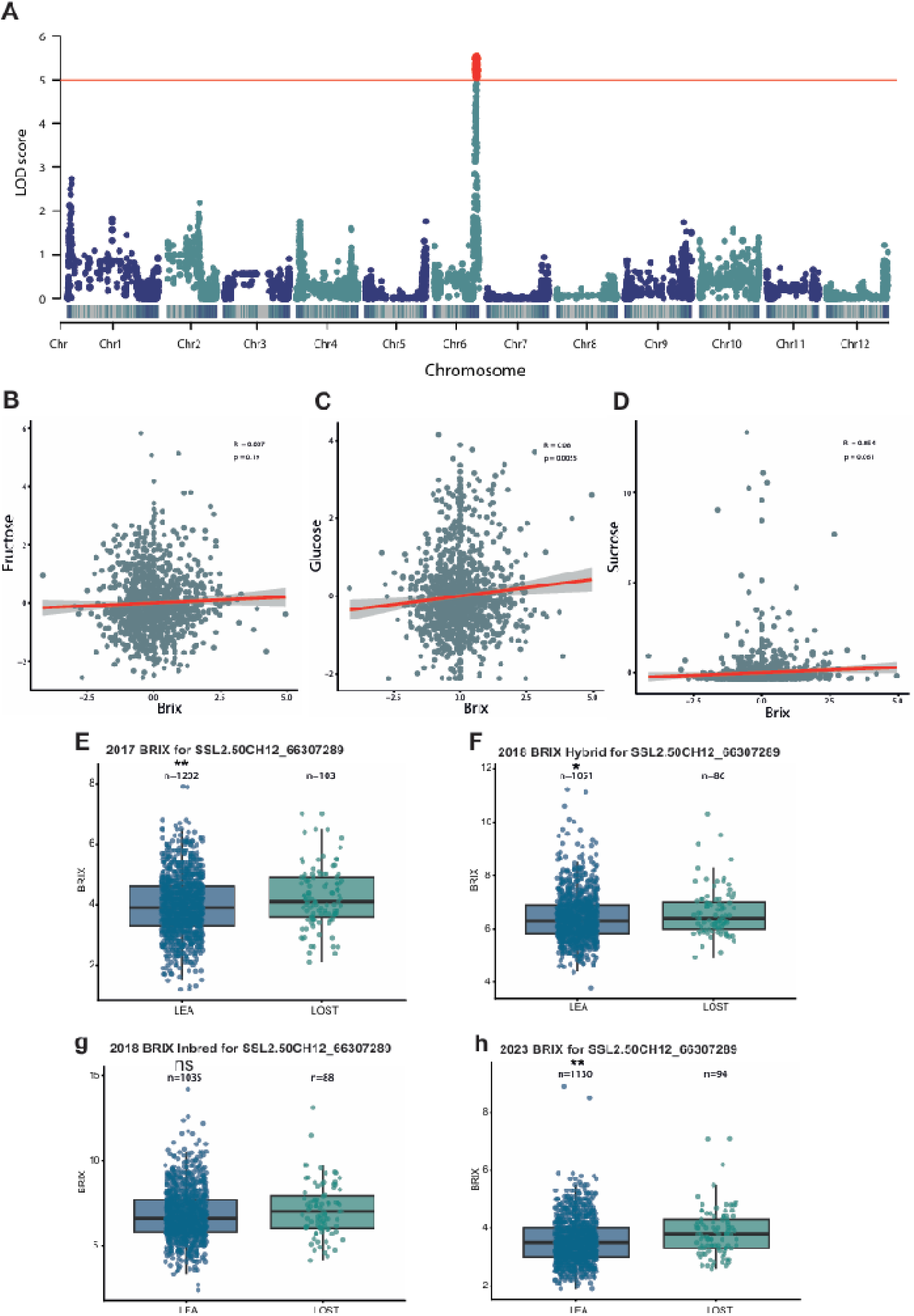
Genetic and metabolic associations with fruit Brix content. **(A)** Brix QTL. **(B)** Correlation between fructose content and Brix. **(C)** Correlation between glucose and Brix. **(D)** Correlation between sucrose and Brix. Brix data was provided by Dani Zamir. Pearson correlation was performed with Z-score normalization of Brix data and metabolite data. **(E, F, G and H)** Boxplots of BRIX measurements for SNP SSL2.50CH12_66307289, corresponding to the peak sucrose-associated locus harboring SlINVINH3 across four datasets: 2017, 2018 hybrid, 2018 inbred and 2023. Distributions are shown separately by allele. Boxplots were generated using the raw phenotypic data. The number of lines representing each allele is indicated within the respective boxes. Statistical significance between allelic classes was asses using Student’s t-test. *<0.05, **<0.01, ***<0.001, ns= not significance.

**Supplemental Figure 7:**
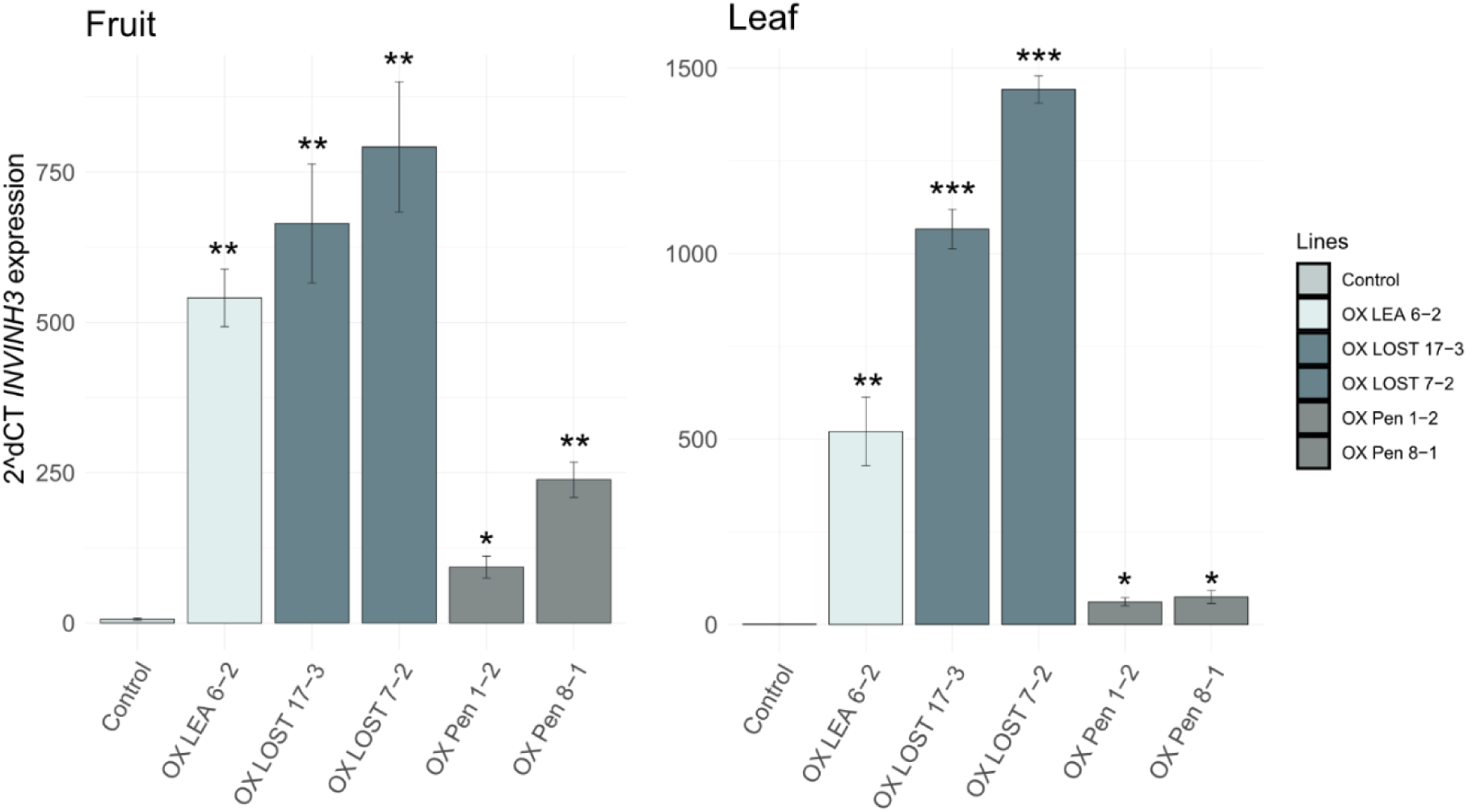
INVINH3 overexpression levels in fruit and leaf tissues. Quantitative RT–PCR analysis of *INVINH3* expression in T1 overexpression (Zhu et al.) lines in fruit and leaf tissues compared with control plants. Bar plots show mean expression levels, with error bars representing standard deviation. Expression was normalized to actin as the housekeeping gene and is shown relative to the control. Statistical significance was determined by comparison with control plants. *<0.05, **<0.01, ***<0.001

**Supplemental Figure 8:**
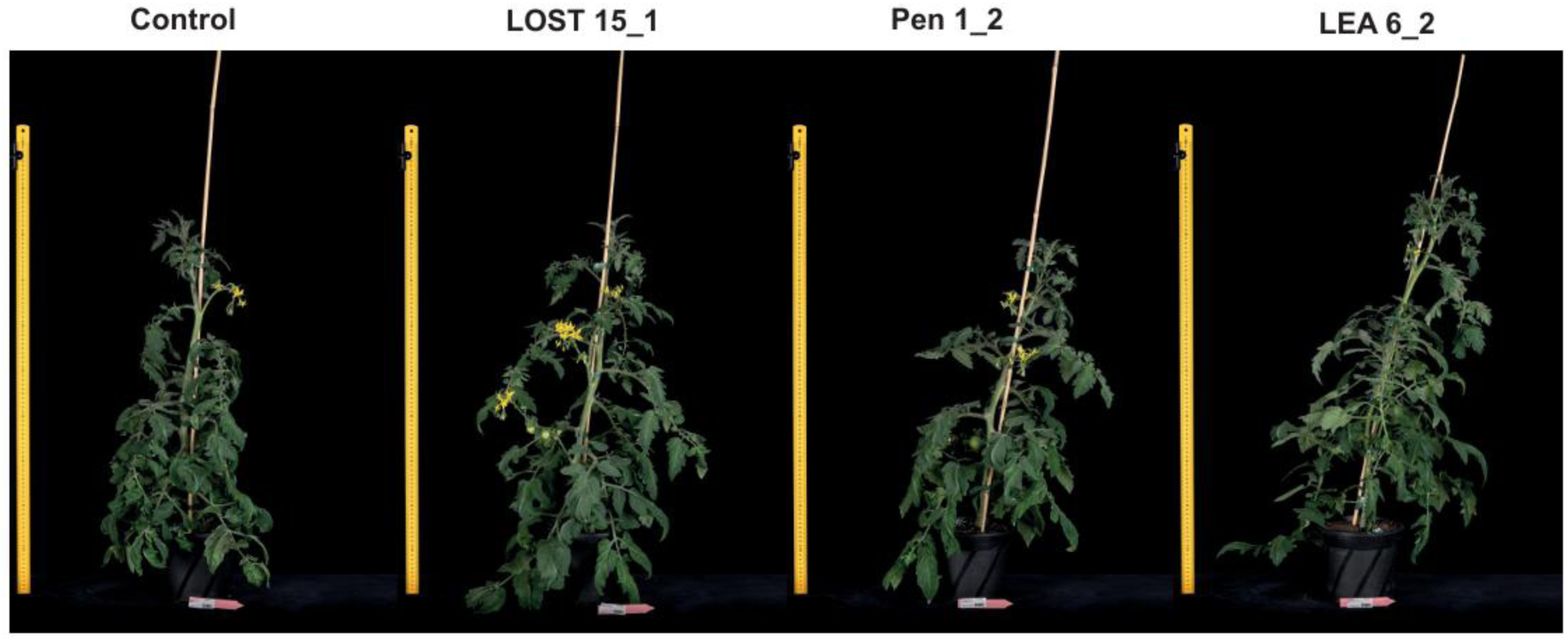
T0 phenotypes of SlINVH3 lines for control (Moneymaker), LOST (*S. pennellii* 5240), Pen (*S.pennellii* 0716) and Lea (domesticated allele).

**Supplemental Figure 9:**
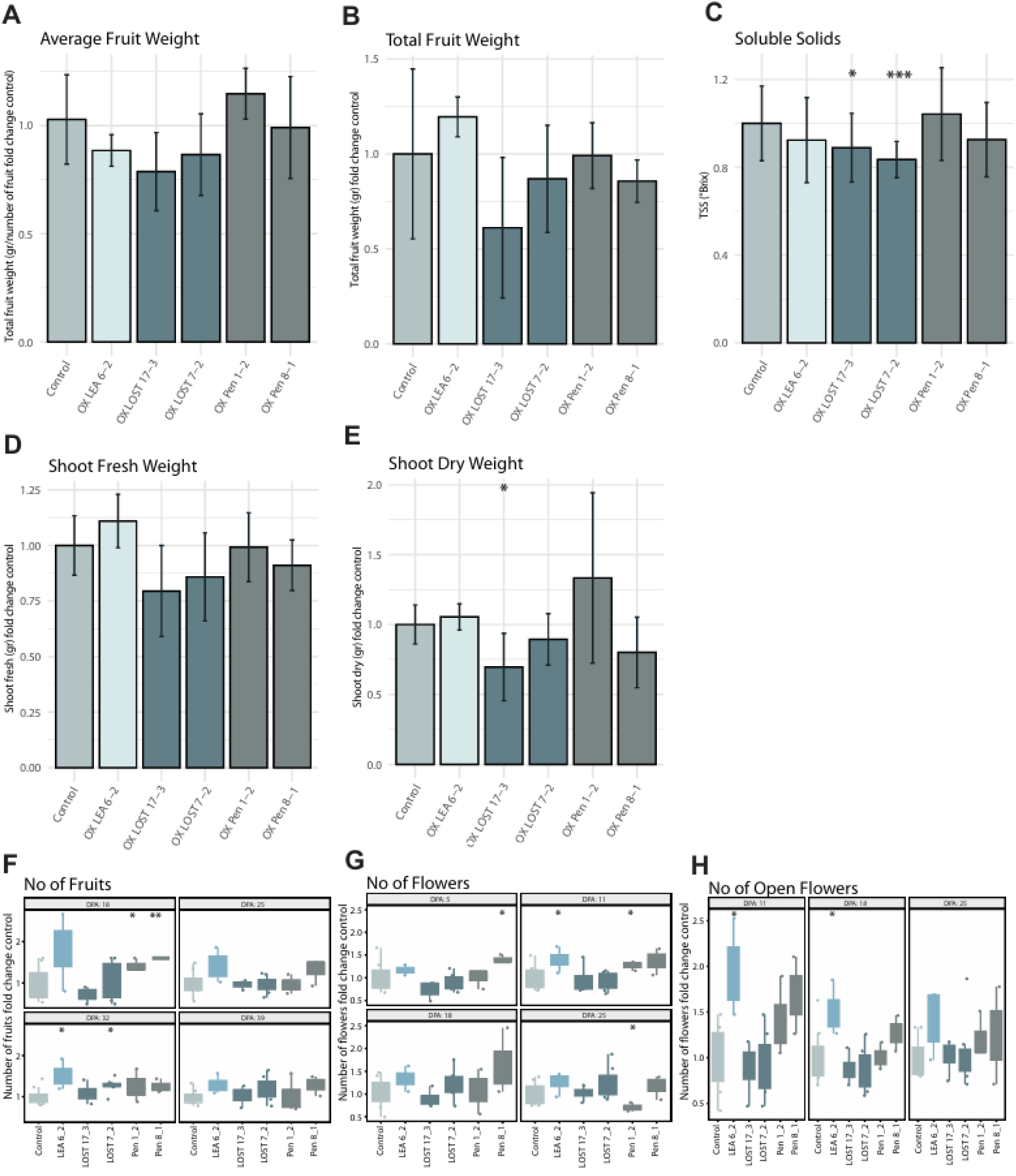
Phenotypic effects of INVINH3 overexpression in T1 lines carrying domesticated and wild alleles. Fold changes in key phenotypic traits of T1 *INVINH3* overexpression lines relative to control ‘Moneymaker’ plants. The analyzed genotypes include LEA 6-2 (domesticated allele), LOST 17-3 and LOST 7-2 (wild-type allele from *S. pennellii* 5240), and Pen 1-2 and Pen 8-1 (wild-type allele from *S. pennellii* 0716). **(A)** Average fruit weight of T1 INVINH OX lines compared with control (money maker) plant; **(B)** Total fruit weight; **(C)** Brix (Total soluble solid contents); **(D)** Shoot fresh weight; **(E)** Shoot dry weight; **(F)** Number of fruits; **(G)** Number of flowers; **(H)** Number of open flowers. p-value *<0.05, ** < 0.01, *** < 0.001.

