## Supplemental Figures for "High-resolution mapping and epistatic QTL of tomato fruit metabolism"

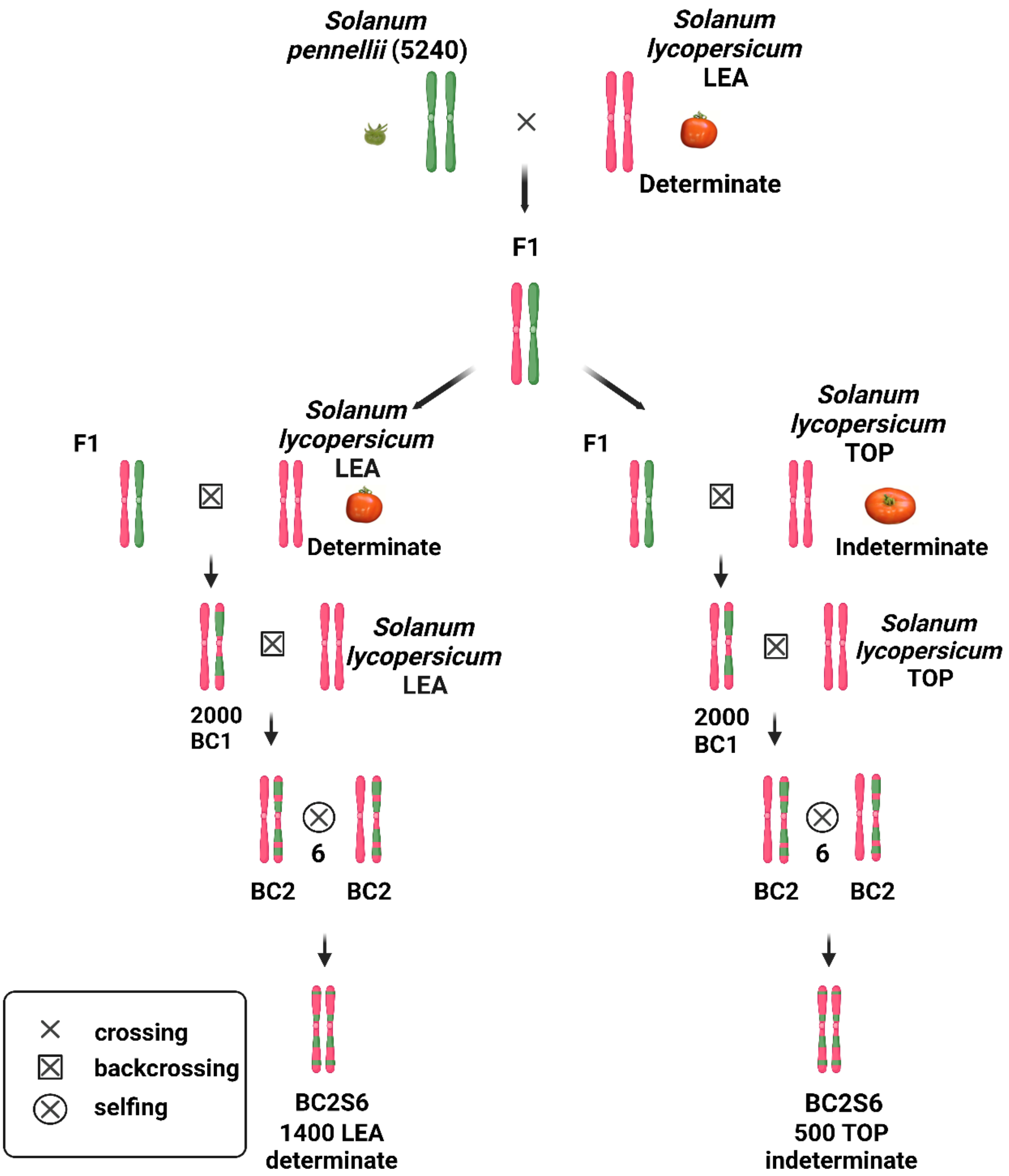


**Supplemental Figure 1:** Breeding scheme of both LEA and TOP backcross inbred lines (BILs) populations. Note that a single plant of the hybrid of LEA and LA5240 gave rise to both the LEA and TOP BILs.

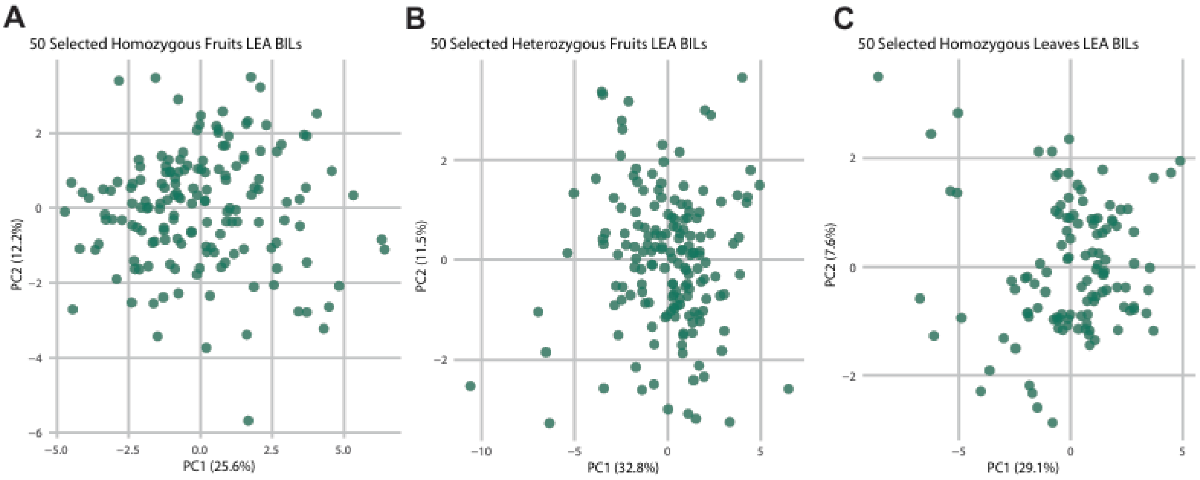


**Supplemental Figure 2: Principal component analysis (PCA) of metabolite profiles from 50 selected LEA BILs. (A, B)** PCA of 50 selected LEA BILs homozygous and heterozygous for fruit. PC1 and PC2 explain 25.6% and 12.2% of the variance for homozygous fruits, and 32.8% and 11.5% for heterozygous fruits. **(C)** PCA of 50 selected LEA BILs homozygous leaves with PC1 and PC2 explaining 29.1% and 7.6% of the variance. Data were log10-transformed, and scaled by the square root of the standard deviation for each variable after internal standard normalization.

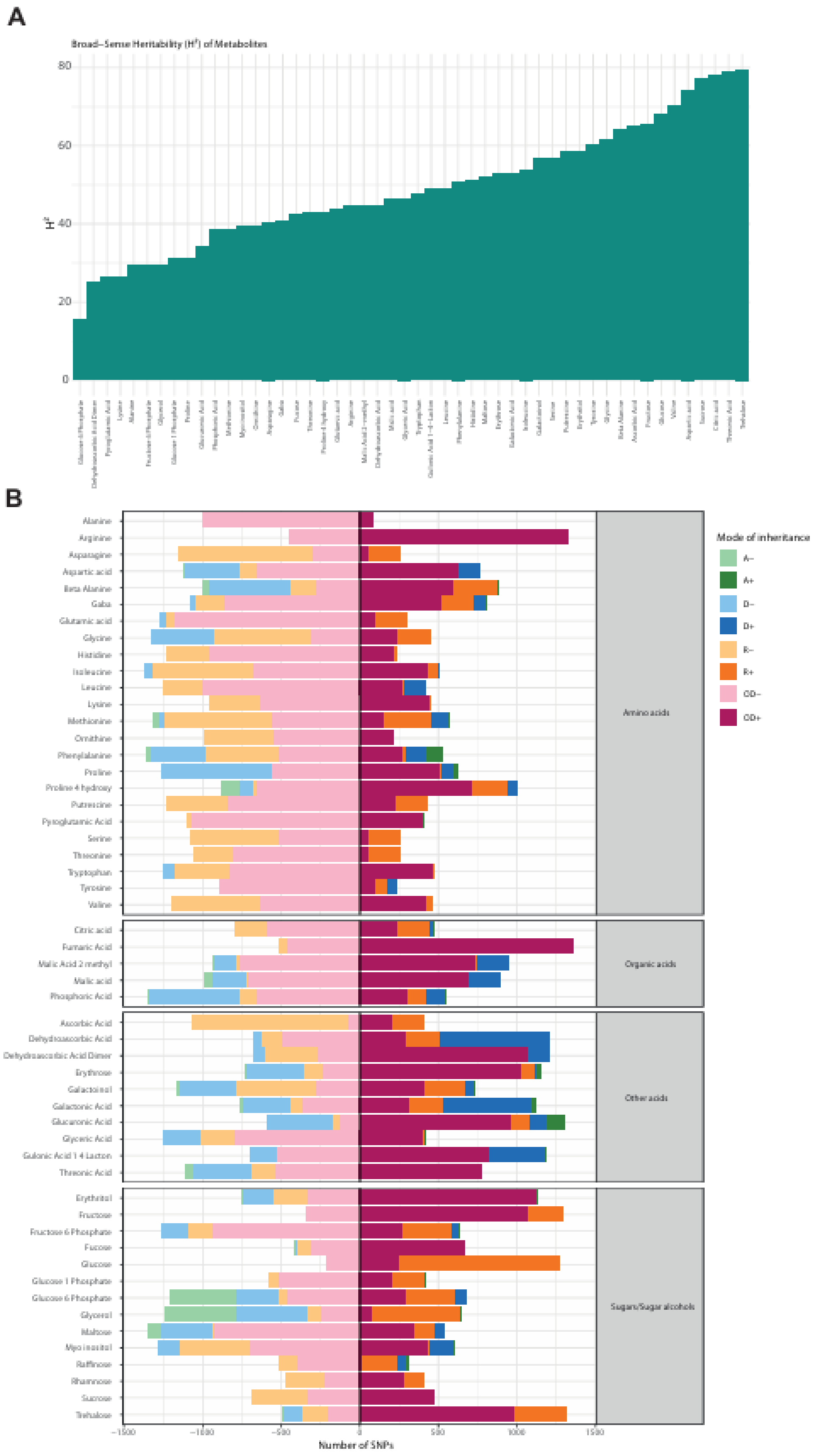


**Supplemental Figure 3: Mode of Inheritance and Broad-Sense Heritability of Metabolites in 50 Selected LEA BIL Population. (A)** SNP-level modes of inheritance were determined across 27 tomato lines using 50 selected LEA BILs, three replicates per homozygous and heterozygous genotype. Marker effects were estimated at the population level to classify allele action as additive (A+ and A−), dominant (D+ and D−), recessive (R+ and R−), or overdominant (OD+ and OD−). Positive signs (+) indicate that the wild allele increases the trait value, whereas negative signs (−) indicate a decrease. The x-axis shows the number of SNPs assigned to each inheritance category. The vertical line separates decreasing (left) and increasing (right) effects of the wild allele for metabolites. **(B)** Broad-sense heritability ($H^2$) of metabolites was estimated using data from 50 LEA backcross inbred lines (BILs) evaluated in a field trial with three biological replicates per line. Variance components were obtained from a random-effects ANOVA model. Heritability estimates ranged from 15% for glucose 6 phosphate (lowest) to 80% for trehalose (highest), indicating substantial variation in the genetic contribution to metabolite accumulation across traits.

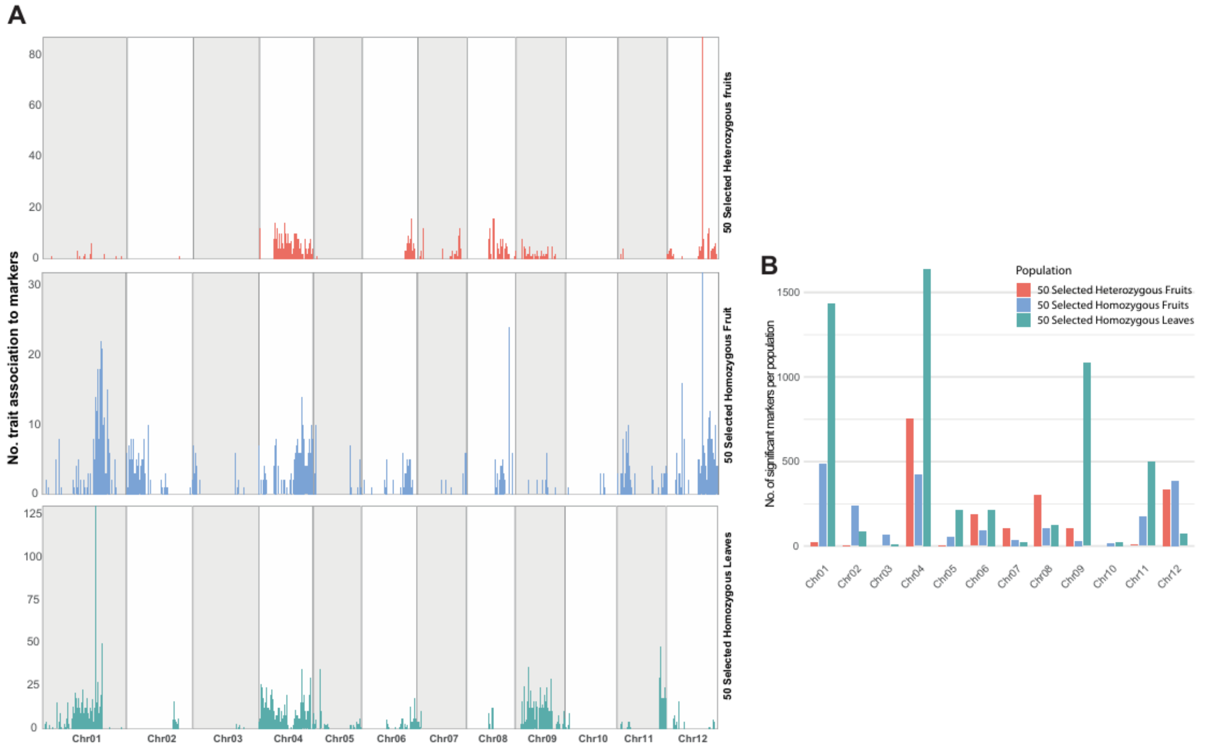


**Supplemental Figure 4: Distribution of QTL-associated markers across chromosomes in the 50 selected LEA BIL populations.** (**A)** Bar charts showing the number of traits associated with significant QTL markers on each chromosome in 50 selected LEA BILs, analyzed separately for heterozygous fruits, homozygous fruits, and homozygous leaves. **(B)** Bar plot showing the total number of significant markers per chromosome in the 50 selected LEA BIL population, combining heterozygous and homozygous fruits and homozygous leaves.

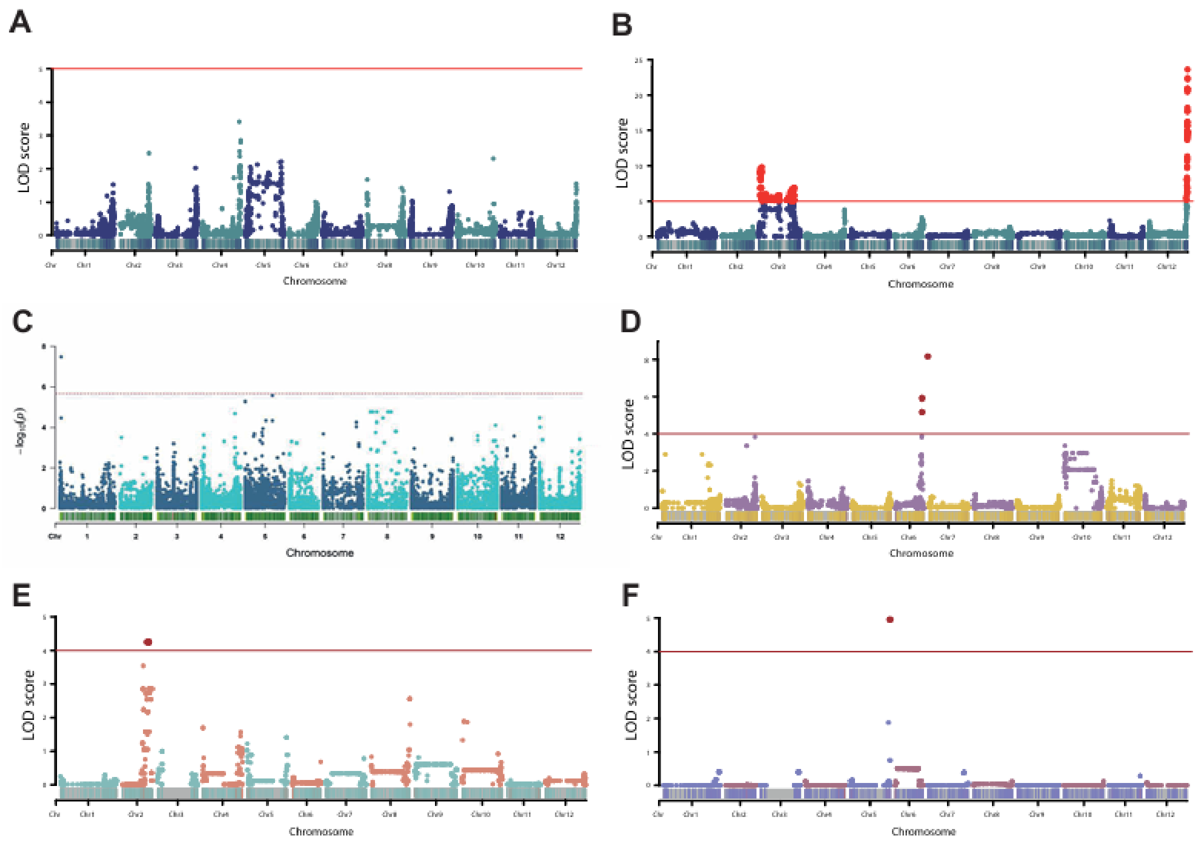


**Supplemental Figure 5: Genome-wide association of sucrose content across tomato populations**. **(A)** 500 TOP BILs 2018 (heterozygous). **(B)** 500 TOP BILs 2019 (homozygous) where the significant QTL is located on chromosome 12. **(C)** 150 lines of GWAS Bulgarian population (unpublished). **(D)** *S. pennellii* (0716) 446 BIL population (unpublished). **(E)** *S. neorickii* 107 BILs homozygous. **(F)** *S. neorickii* 107 BILs heterozygous (Brog et al., 2019).

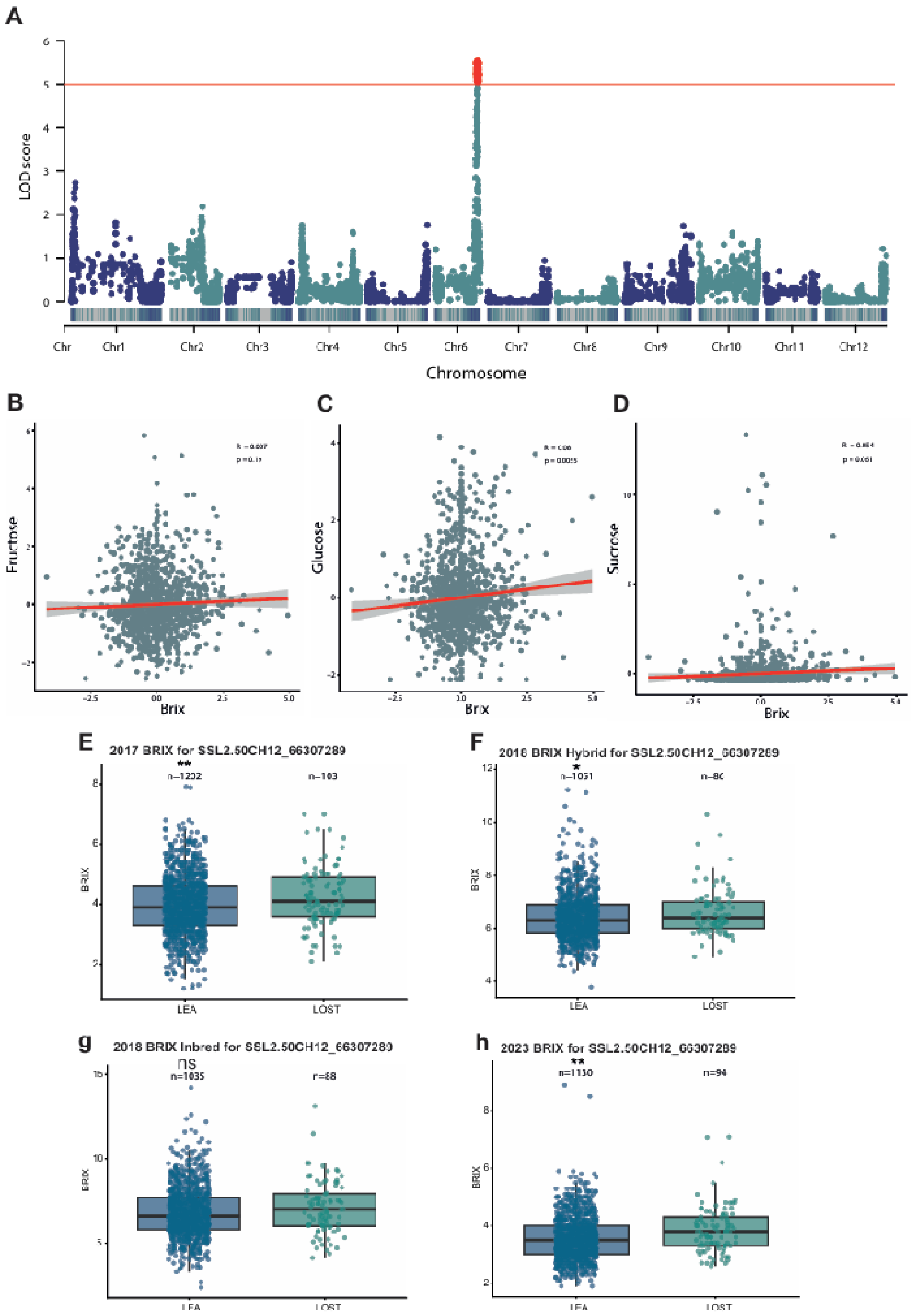


**Supplemental Figure 6: Genetic and metabolic associations with fruit Brix content. (A)** Brix QTL. **(B)** Correlation between fructose content and Brix. **(C)** Correlation between glucose and Brix. **(D)** Correlation between sucrose and Brix. Brix data was provided by Dani Zamir.

Pearson correlation was performed with Z-score normalization of Brix data and metabolite data. **(E, F, G and H)** Boxplots of BRIX measurements for SNP SSL2.50CH12_66307289, corresponding to the peak sucrose-associated locus harboring SlINVINH3 across four datasets: 2017, 2018 hybrid, 2018 inbred and 2023. Distributions are shown separately by allele. Boxplots were generated using the raw phenotypic data. The number of lines representing each allele is indicated within the respective boxes. Statistical significance between allelic classes was asses using Student's t-test. *<0.05, **<0.01, ***<0.001, ns= not significance.

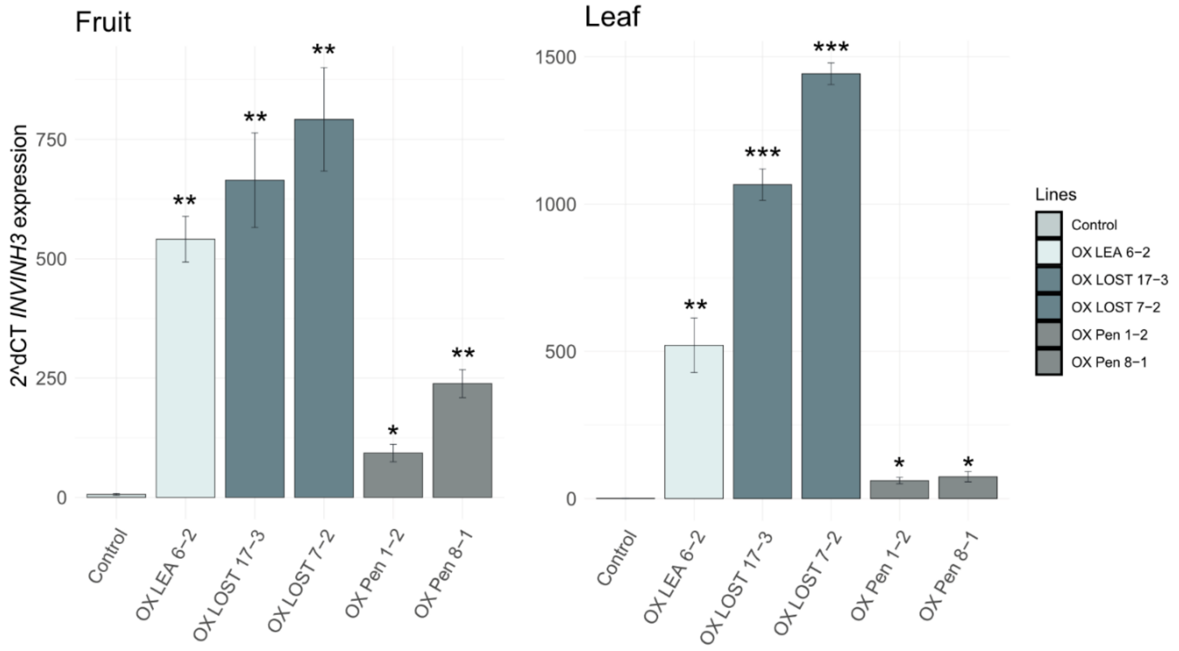


**Supplemental Figure 7: INVINH3 overexpression levels in fruit and leaf tissues**. Quantitative RT–PCR analysis of *INVINH3* expression in T1 overexpression (Zhu et al.) lines in fruit and leaf tissues compared with control plants. Bar plots show mean expression levels, with error bars representing standard deviation. Expression was normalized to actin as the housekeeping gene and is shown relative to the control. Statistical significance was determined by comparison with control plants. *<0.05, **<0.01, ***<0.001

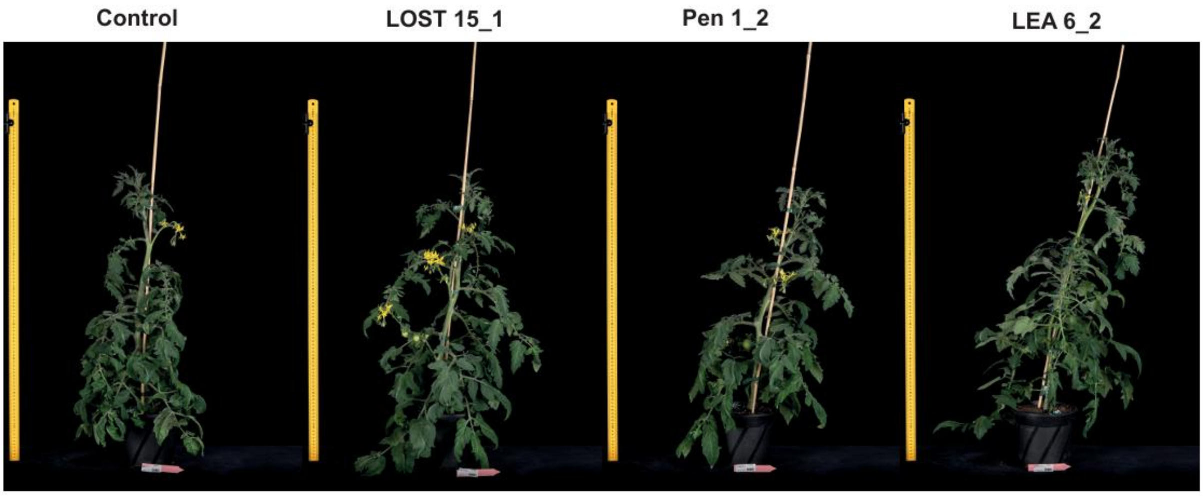


**Supplemental Figure 8:** T0 phenotypes of SlINVH3 lines for control (Moneymaker), LOST (*S. pennellii* 5240), Pen (*S.pennellii* 0716) and Lea (domesticated allele).

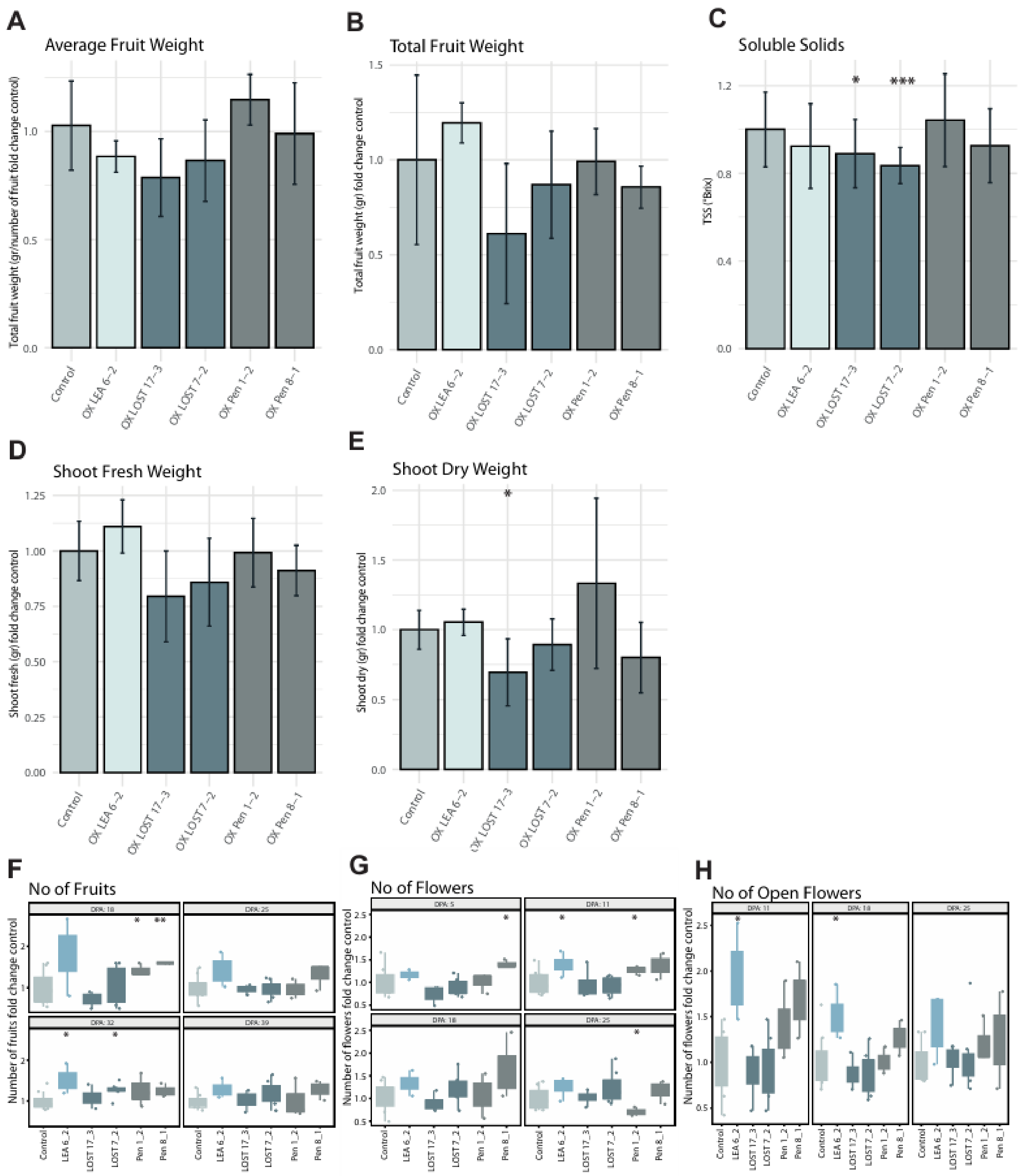


**Supplemental Figure 9: Phenotypic effects of INVINH3 overexpression in T1 lines carrying domesticated and wild alleles.** Fold changes in key phenotypic traits of T1 *INVINH3* overexpression lines relative to control 'Moneymaker' plants. The analyzed genotypes include LEA 6-2 (domesticated allele), LOST 17-3 and LOST 7-2 (wild-type allele from *S. pennellii* 5240), and Pen 1-2 and Pen 8-1 (wild-type allele from *S. pennellii* 0716). **(A)** Average fruit weight of T1 INVINH OX lines compared with control (money maker) plant; **(B)** Total fruit weight; **(C)** Brix (Total soluble solid contents); **(D)** Shoot fresh weight; **(E)** Shoot dry weight; **(F)** Number of fruits; **(G)** Number of flowers; **(H)** Number of open flowers. p-value *<0.05, ** < 0.01, *** < 0.001.
